# Structural Basis for Cas9-Directed Spacer Acquisition in Type II-A CRISPR-Cas Systems

**DOI:** 10.1101/2025.06.12.659315

**Authors:** Zhaoxing Li, Yutao Li, Jianping Kong, Qianqian Wu, Pingping Huang, Yu Zhang, Wanqian Wu, Meirong Chen, Yongfeng Liu, HanFeng Lin, Liqiu Hou, Gongyu Liu, Ting Zeng, Yutong He, Chunyi Hu, Zhenhuang Yang, Meiling Lu, Min Luo, Yibei Xiao

## Abstract

CRISPR-Cas systems confer prokaryotic adaptive immunity by integrating foreign DNA (prespacers) into host arrays. Type II-A systems employ Cas9 for protospacer adjacent motif (PAM) recognition and coordinate with Csn2 and the Cas1-Cas2 integrase during spacer acquisition, yet their structural basis remains unresolved. Here, we report cryo-EM structures of the *Enterococcus faecalis* Cas9-Csn2-Cas1-Cas2 supercomplex in apo and DNA-bound states. The apo-state structure (Cas9_2_-Csn2_8_-Cas1_8_-Cas2_4_) adopts a resting conformation, with Cas9 locked in a nuclease-inactive state and Cas1-Cas2 sterically blocked from prespacer loading. Upon DNA engagement, Cas9 undergoes a conformational transition, forming a prespacer catching complex that threads the DNA through Csn2’s central channel. This architecture enables Cas9 to interrogate the PAM sequence while sliding along the DNA, with Cas9 and Csn2 jointly define a 30-bp DNA segment which matches the prespacer length. Subsequent dissociation of Cas9 triggers a structural reconfiguration of the Csn2-Cas1-Cas2 assembly. The PAM-proximal DNA becomes accessible, and Cas1-Cas2 relocates to bind to the exposed DNA, enabling further prespacer processing and directional integration. These findings reveal how Cas9 collaborates with Csn2 and Cas1-Cas2 to couple PAM recognition with prespacer selection, resolving the dynamic structural transitions that ensure fidelity during type II-A CRISPR adaptation.

## INTRODUCTION

Clustered Regularly Interspaced Short Palindromic Repeats (CRISPR) and CRISPR associated proteins (Cas) endow prokaryotes with an RNA-guided adaptive immunity.^1–4^ Among these systems, the RNA-guided interference activity of type II-A CRISPR-Cas9 has been extensively utilized for genome editing.^5^ CRISPR-Cas systems establish immunity by inserting short foreign DNA fragments (prespacer) into CRISPR arrays in a process termed spacer acquisition.^6^ These arrays subsequently transcribe CRISPR-derived RNAs (crRNAs) that direct Cas effectors to neutralize invasive genetic elements through sequence-specific interference. While CRISPR systems exhibit diversity in their interference machinery, they all rely on the conserved Cas1-Cas2 integrase for spacer acquisition.^7–12^

In DNA-targeting CRISPR systems, the Cas1-Cas2 integrase complex acquire spacers preferentially from foreign DNA harboring a 3–5 base pair (bp) protospacer-adjacent motif (PAM), a critical mechanism to prevent self-targeting.^7^ Recent studies have elucidated how PAM-containing prespacers are selected, processed and integrated step wisely to establish directional integration relative to the PAM position in certain systems.^13–23^ The Cas1-Cas2 hexamer preferentially loads partially duplexed DNA with 3’ single-stranded overhangs bearing a PAM sequence.^7^ In type I-E and I-F systems, host nucleases such as DnaQ exonuclease process the non-PAM overhangs, while Cas1 shields the PAM-proximal overhangs. In other systems (e.g., type I, II, and V), Cas4 mediates PAM protection and processing.^13–23^ Following leader-proximal integration of the processed non-PAM overhang, DnaQ or Cas4 cleaves the PAM-proximal overhang, releasing processing factors and enabling full integration by Cas1-Cas2.^18, 24–28^ These findings highlight how PAM sequestration and asymmetric overhang trimming coordinate prespacer integration orientation. However, the prespacer selection mechanism in type II-A CRISPR-Cas9 systems remains unresolved, as Cas1 lacks intrinsic PAM recognition and Cas4 is absent.

Type II-A systems encode four proteins: the Cas9 effector nuclease, Cas1-Cas2 integrase, and the auxiliary protein Csn2, all essential for in vivo spacer acquisition (Figure 1A).^29,30^ Csn2 forms a tetramer with a central positively charged channel for dsDNA binding, though its precise role is unclear.^31–35^ Previous investigations have captured how the type II-A Cas1-Cas2 integrase complex integrate the trimmed prespacers stepwise *in vitro*.^8,36,37^ Cryo-EM analysis of the type II-A *Streptococcus thermophilus* system revealed a predominant Csn2_8_-Cas1_8_-Cas2_4_ complex, wherein two Cas1-Cas2 hexamers bridge a pair of Csn2 tetramers, forming a channel accommodating approximately 30 bps of duplex DNA.^31^ However, these complexes lack PAM recognition specificity, rendering them unable to specify PAM-containing prespacers. Intriguingly, the effector Cas9 is responsible for PAM recognition in type II-A systems and has been hypothesized to assemble into a Cas9-Csn2-Cas1-Cas2 supercomplex to coordinate spacer acquisition.^29,38,39^ While recent work implicates Cas9’s exonuclease activity in prespacer trimming, this activity is dispensable in vivo, and the molecular basis of PAM specificity remains unknown.^40^ Thus, structural insights are required to unravel the function and architecture of the Cas9-Csn2-Cas1-Cas2 supercomplex in spacer acquisition.

**Figure 1.**
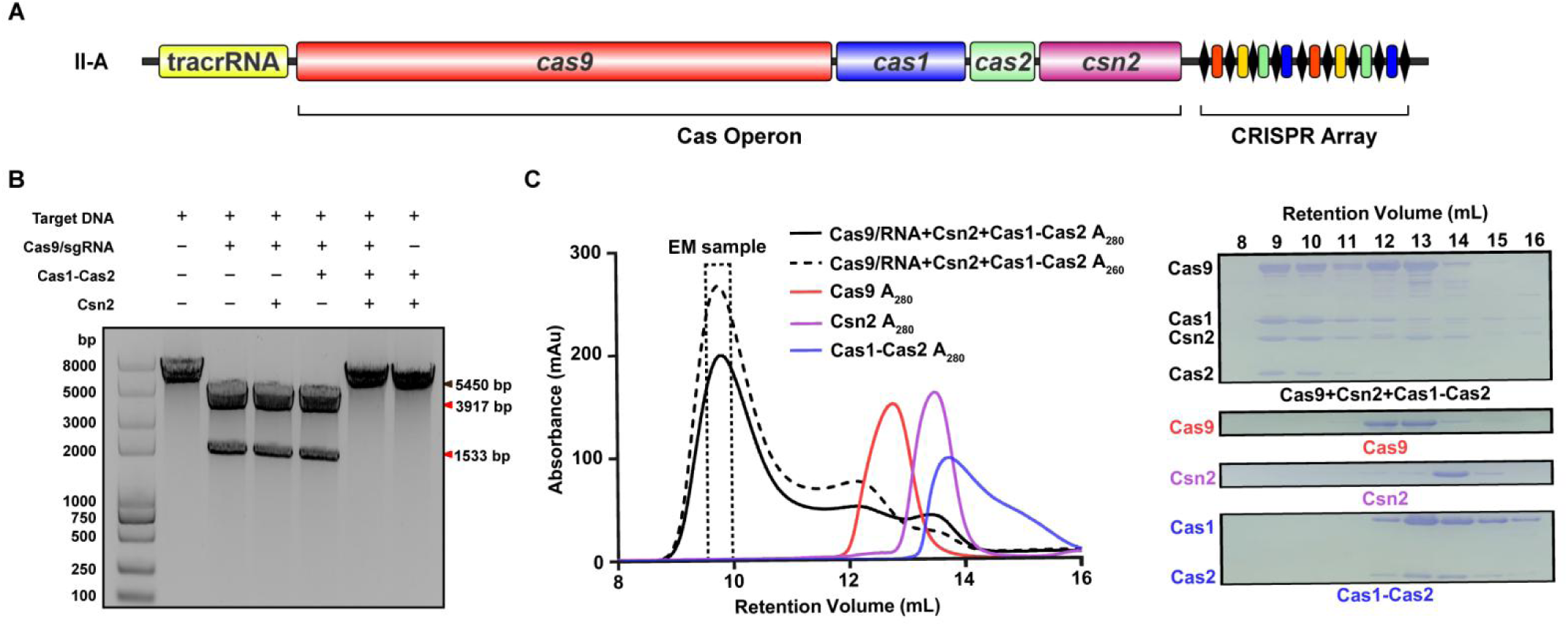
Cas9, Csn2, Cas1, and Cas2 from the *Enterococcus faecalis* type II-A CRISPR-Cas system form a supercomplex in vitro. **(A)** Schematic representation of the type II-A CRISPR-Cas locus of *E. faecalis*. **(B)** *Efa*Cas9 loses its nuclease activity for cleaving target DNA upon forming a supercomplex with Csn2 and Cas1-Cas2. **(C)** Size exclusion chromatography (left) and SDS-PAGE analysis (right) confirmed the formation of the *Efa*Cas9/RNA-Csn2-Cas1-Cas2 supercomplex.

## RESULTS

### Reconstitution and Cryo-EM Structure of the Apo Cas9-Csn2-Cas1-Cas2 Supercomplex

Previous studies demonstrated that the Cas1, Cas2, Csn2 and Cas9 proteins from *Streptococcus thermophilus* and *Streptococcus pyogenes* can form a supercomplex.^29–31,39^ However, obtaining a stable supercomplex remained a challenge and hindered structural characterization. Here, we investigated the type II-A system from *Enterococcus faecalis*. *Efa*Cas9 shares 52% sequence identity with *Spy*Cas9 and also utilizes a 5’-NGG-3’ PAM sequence for target recognition (Figure S1). Notably, we found that the target DNA cleavage activity of Cas9 was markedly inhibited in the presence of both Csn2 and Cas1-Cas2, suggesting supercomplex formation (Figure 1B). Size exclusion chromatography (SEC) confirmed a stable quaternary complex formation (Figure 1C).

To determine the structure, we conducted single-particle cryo-electron microscopy (cryo-EM) analysis on the apo-state Cas9-Csn2-Cas1-Cas2 supercomplex. By carefully analyzing 2-dimensional (2D) and 3-dimensional (3D) reconstructions, we obtained a medium-resolution map of the complex at 3.9 Å resolution (Figure 2A and S2). The overall structure adopts a globular morphology with dimensions of 185 Å in length, 195 Å in width, and 210 Å in thickness. The structure features a distinctive shape with four prominent lobes at the corners, corresponding to four Cas1 dimers (Figure 2B and S3A). The unique orientation of these dimers disrupts the 4-fold symmetry around the central opening. Cas2 dimer density is weakly observed between the Cas1 dimer pairs, however, their C-terminal β strands which interact with the β sheet of the Cas1 N-terminal domain display strong density, confirming the presence of a Cas2 dimer linking two Cas1 dimers (Figure S3A-S3C). Two large rings, representing two Csn2 tetramers, bridge the Cas1-Cas2 hexamers around the central hole. Notably, two substantial blocks of additional density are connected to the upper Cas1-Cas2 hexamer, likely disrupting symmetry and resulting in different Cas1 dimer orientations at the corner. While most density appears weak and devoid of recognizable structural features, each can roughly accommodate one Cas9 monomer, as a result two Cas9 molecules were sandwiched by a pair of Cas1 dimer (Figure 2A and S3A). This is not due to the low occupancy of Cas9 because the REC3 domain that directly interact with Cas1 and Csn2 is well resolved and can be unambiguous modelled (Figure 2B and S3A). Taken together, the apo-state supercomplex structure captured here contains all four proteins with the stoichiometry Cas9_2_-Csn2_8_-Cas1_8_-Cas2_4_, wherein two dumbbell shape Cas1_4_-Cas2_2_ hexamers encircling two Csn2 tetramer rings to encapsulate two Cas9 molecules.

**Figure 2.**
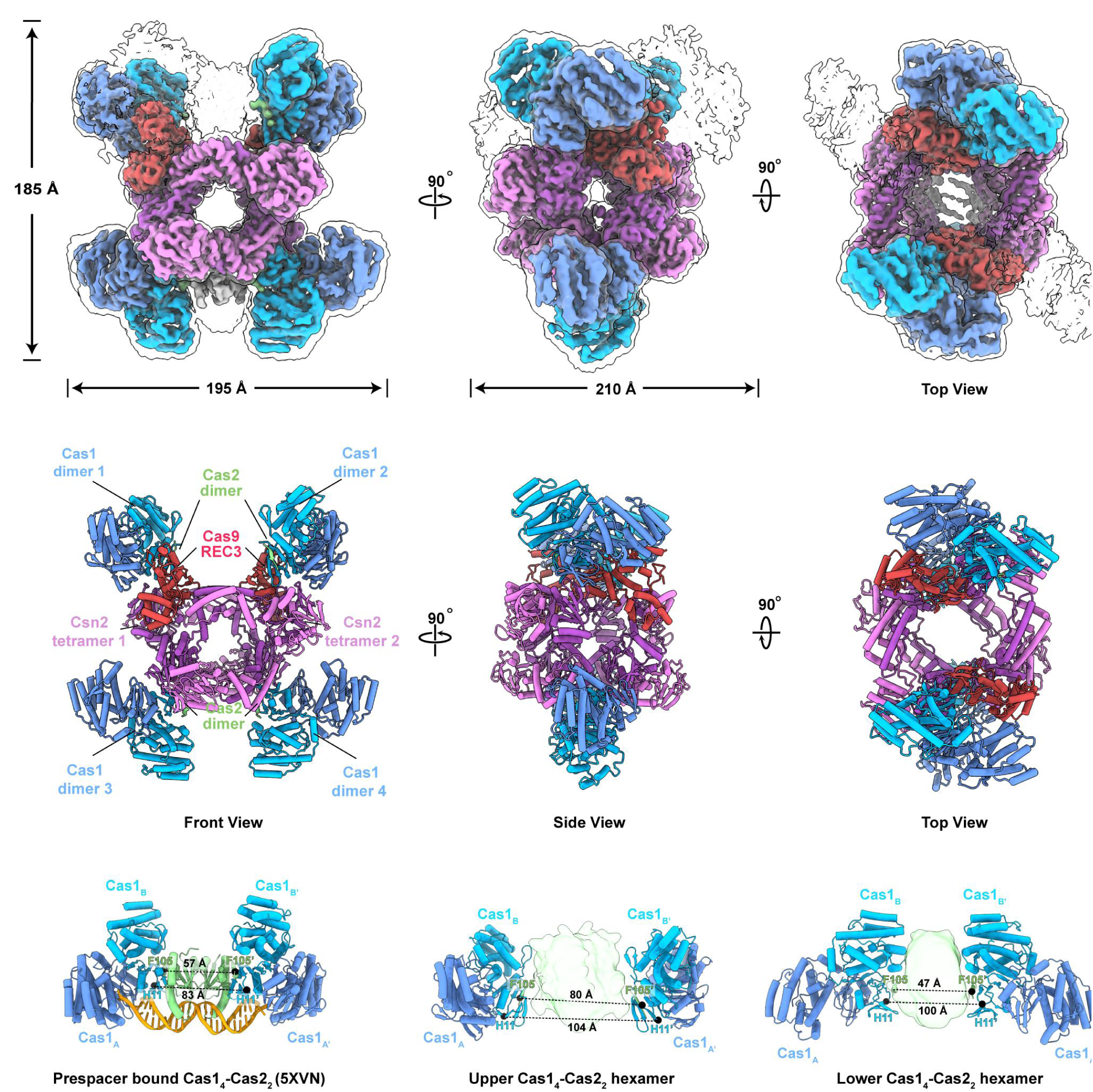
Global and local Cryo-EM structures of apo state Cas9/RNA_2_-Csn2_8_-Cas1_8_-Cas2_4_ supercomplex. **(A)** Cryo-EM density map of the Cas9/RNA_2_-Csn2_8_-Cas1_8_-Cas2_4_ supercomplex. The raw electron density was shown as a transparent map. Subunits are colored as follows using sharpen map filtered to estimated overall resolution: Cas9_REC3_, red; Csn2, purple and violet; Cas1, blue and cyan; Cas2, green. **(B)** The corresponding views of the molecular model. **(C)** Two distinct conformations of the Cas1_4_–Cas2_2_ hexamer within the supercomplex, compared with the previously resolved protospacer-bound Cas1–Cas2 structure (PDB 5XVN), reveal its conformational expansion and contraction in the Cas9/RNA_2_-Csn2_8_-Cas1_8_-Cas2_4_ complex, with distances measured between F105 of each Cas2 and His11 of each catalytic Cas1.

### Apo-state Cas9-Csn2-Cas1-Cas2 supercomplex represent a resting conformation

While the REC3 domain of Cas9 is well-resolved, the remaining domains exhibit pronounced flexibility. Despite this, the overall architecture is discernible, enabling comparative docking studies with previously determined Cas9 structures in diverse functional states. Attempts to align existing *Spy*Cas9 structures (e.g., apo-state (PDB 4CMP) or closed-state (PDB 7S36)) with the density map were unsuccessful (Figure S3D and S3E), indicating that Cas9 adopts a novel conformation within the supercomplex. This unique configuration is stabilized by extensive interactions between the REC3 domain and both Cas1 and Csn2 (Figure 2B). The REC3 domain is known to allosterically regulate the HNH nuclease domain, undergoing substantial conformational rearrangements upon target DNA binding.^41^ Notably, REC3 deletion abolishes Cas9’s cleavage activity while preserving DNA-binding affinity, underscoring its role in modulating nuclease function.^42–45^ The observed REC3 immobilization within the supercomplex likely restricts HNH domain activation, consistent with the attenuated DNA cleavage activity detected in our assays (Figure 1B). By locking Cas9 in a REC3-constrained state, the supercomplex may enable DNA surveillance without promiscuous cleavage, a critical safeguard for maintaining precision during spacer acquisition.

The four Cas1 dimers and two Csn2 tetramers in the supercomplex adopt conformations analogous to those reported in prior structural studies.^8,35^ The N-terminal domains of Cas1 dimers engage the head regions of Csn2 tetramers, recapitulating interfaces observed in the *St*Csn2-dsDNA-Cas1-Cas2 monomer (PDB 6QXF). However, significant structural deviations arise in the Csn2-Cas1-Cas2 assembly. Unlike the parallel alignment of Csn2 tetramers along the DNA duplex in the *St*Csn2-dsDNA-Cas1-Cas2 complex, the two Csn2 tetramers here tilted by 38°, forming a V-shaped architecture (Figure S4A). Tilting of the Csn2 tetramers induces contraction of the lower Cas1-Cas2 hexamer and expansion of the upper hexamer, creating an enlarged cavity to accommodate two Cas9 molecules (Figure 2B). Consequently, the two Cas1-Cas2 hexamers adopt asymmetric configurations, with Cas1 dimers exhibiting clockwise or counterclockwise rotations relative to the prespacer-bound Cas1-Cas2 structure (PDB 5XVN) (Figure S4B).

The Cas2 C-terminal tails further reflect structural plasticity. The inter-tail distances measure 80 Å and 47 Å for the upper and lower hexamer, respectively, diverging markedly from the 57 Å spacing in prespacer-bound Cas1-Cas2 (Figure 2C). This flexibility suggests Cas2 C-terminal tail dynamics regulate hexamer conformational states. Cas1-Cas2 hexamer specify a splayed prespacer containing a 22-bp duplex and two 4-nt 3’-overhangs by end-stacking of two His11 in each catalytic Cas1.^8^ However, the distance between catalytic His11 residues in the upper and lower Cas1-Cas2 hexamers are 104 Å and 100 Å, respectively, exceeds the 83 Å distance observed in prespacer-bound structures (PDB 5XVN), indicating a relaxed configuration incompatible with prespacer loading. Critically, steric occlusion by the Csn2 tetramers blocks DNA access to both hexamers, precluding prespacer loading for integration.

Collectively, these structural features define the apo supercomplex as a resting state. This conformation enables Cas9 to engage DNA for target interrogation while suppressing nuclease activity, thereby minimizing off-target cleavage. Simultaneously, the steric and conformational constraints imposed on Cas1-Cas2 hexamers prevent premature prespacer loading, reducing the risk of aberrant spacer integration. Such regulatory mechanisms likely ensure fidelity and efficiency during CRISPR adaptive immunity, balancing DNA surveillance with stringent control over cleavage and integration processes.

### PAM-containing DNA selection by the prespacer catching complex

To elucidate the molecular mechanism underlying selective recognition of PAM-containing DNA by the Cas9-Csn2-Cas1-Cas2 supercomplex, we designed a 59-bp dsDNA substrate devoid of crRNA complementarity but harboring a single 5’-NGG-3’ PAM. This DNA substrate was incubated with the supercomplex at a molar ratio of 1.2:1, followed by cryo-electron microscopy (cryo-EM) data collection. In contrast to the homogeneous apo-state Cas9-Csn2-Cas1-Cas2 supercomplex, the DNA-bound complex exhibited significant morphological heterogeneity, indicating substantial structural rearrangements upon DNA binding. Through particle selection and subsequent 2D and 3D classification, We obtained reconstructions of two relatively intact Cas9-containing complexes, referred to as the prespacer catching complex, with overall resolutions of 2.96Å (State I, EMD-61100) and 3.01 Å (State II, EMD-61101), respectively. (Figure S5).

The dimensions of the prespacer catching complex are approximately 178 Å × 151 Å × 98 Å, which is less than half the size of the apo supercomplex (Figure 3A). The resulting atomic model consists of one Cas9/crRNA/tracrRNA subunit, one Csn2 tetramer, one Cas1 dimer, and a dsDNA strand threaded through the Csn2 central channel, binding to Cas9 at both the PAM-interacting domain and the REC2 domain (Figure 3B). Notably, no distinct density for the Cas2 dimer or the second Cas1 dimer was observed, suggesting a high degree of flexibility of these molecules within the prespacer catching complex. Unlike the highly flexible Cas9 in the apo-state supercomplex, Cas9 within the prespacer catching complex exhibits a much more stable conformation, thereby allowing for a clear distinction of various domains (Figure S6). Cas9’ conformational arrangement upon DNA binding is the driving force that leads to the disassembly of the apo supercomplex. The REC and NUC lobes of Cas9 adopt a closed conformation that effectively clamps the DNA duplex, inducing a pronounced bend and underwinding of the DNA at the candidate complementarity region (CCR) (Figure 3C). Cas9 structures in the two states exhibit distinct conformational changes within the REC2 domain (Figure S6D). The State I Cas9 structure closely resembles the SpyCas9/sgRNA/DNA complex with a 3 bp R-loop (PDB: 7S38) (Figure S6E), whereas the State II structure aligns more closely with SpyCas9/sgRNA/DNA lacking RNA : DNA base pairs (PDB: 7S36) (Figure S6F). This indicates that both states represent DNA interrogation intermediates.

**Figure 3.**
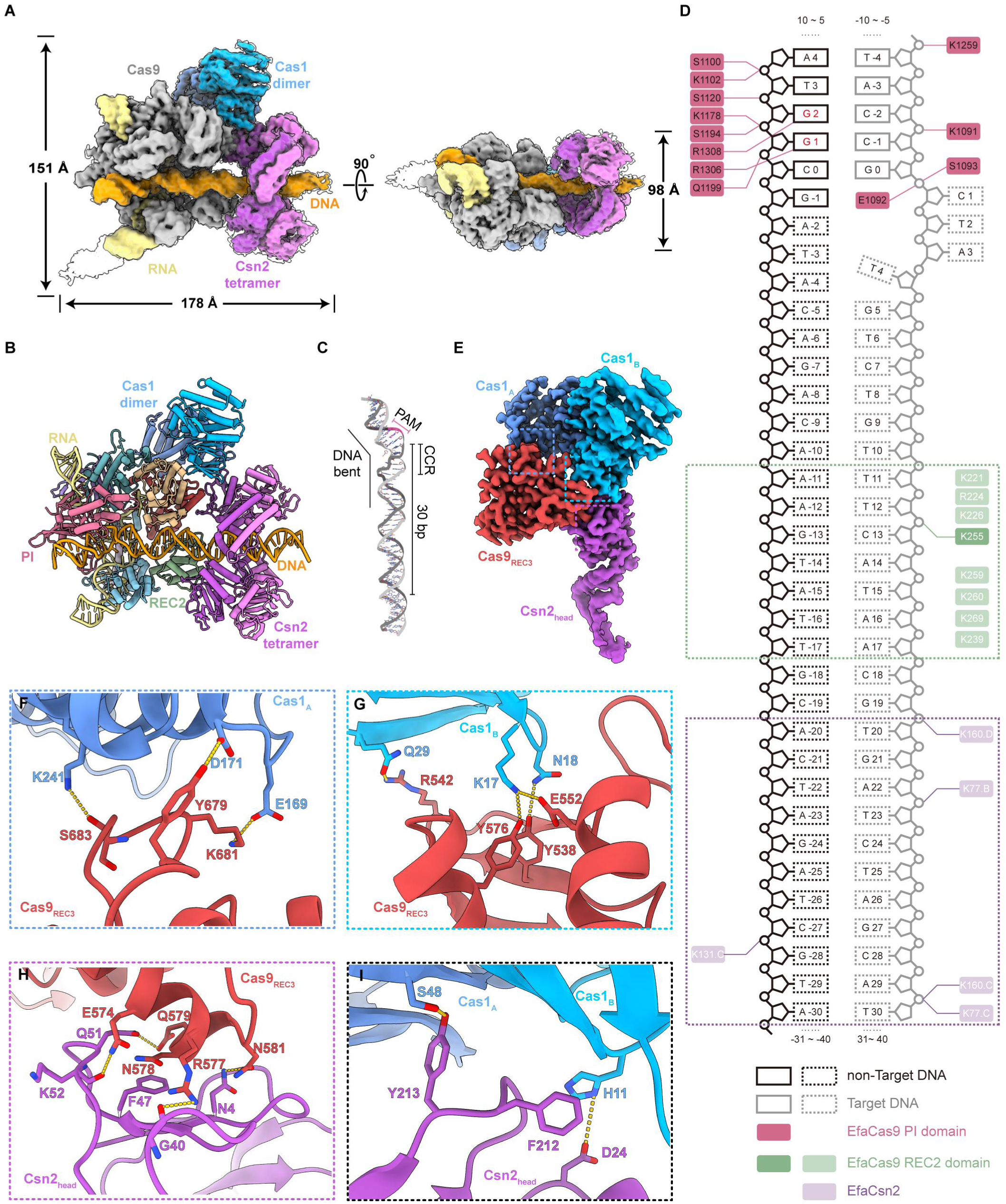
Rearrangement of Cas9, Csn2, and Cas1 in the Cryo-EM structures of the prespacer capture complex. **(A)** Cryo-EM density map of the Cas9/RNA-Csn2-Cas1-DNA prespacer capture complex, state I. The raw electron density was shown as a transparent map. Subunits are colored as follows: Cas9, sliver; Csn2, purple and violet; Cas1, blue and cyan; DNA, orange; RNA, Khaki. Colored map were filtered by Guassian with width set to 1 in Chimera X. **(B)** The molecular model of the prespacer capture complex with each domain of Cas9 well resolved. **(C)** DNA bending and underwinding at the candidate complementarity region. **(D)** Schematic illustration of the molecular interactions between DNA and Cas proteins. Black boxes represent non-Target DNA. Grey boxes represent Target DNA. Pink boxes are assigned to the EfaCas9 PI domain. Green boxes are assigned to the EfaCas9 REC2 domain. Purple boxes are assigned to EfaCsn2. Solid-line borders and dark hues denote regions exhibiting well-resolved electron density. Dashed borders and light hues denote regions exhibiting poor density. **(E-I)** The REC3 domain of Cas9 interacts with Cas1 and Csn2 in the prespacer capture complex. Hydrogen bonds are shown as yellow dash.

Cas9 binds DNA in a PAM-dependent manner, with the residues R1306 and R1308 specifically recognizing the guanine nucleobases dG1 and dG2 of the PAM sequence in the non-target strand through base-specific hydrogen-bonding interactions (Figure 3D). The PAM-interacting domain of Cas9 further engages with the minor groove of the PAM duplex, where a cluster of positively charged residues (K1091/K1102/K1178/K1259) and serine residues (S1100/S1120/S1194) interact with the negatively charged DNA phosphate backbone, stabilizing DNA binding (Figure 3E). Residues E1092 and S1093 of the “phosphate lock loop” interact with the phosphodiester group linking dG0 and dC1 in the target DNA strand, contributing to the flipping of the target DNA strand at the bent CCR. This bend-induced base-flipping mechanism has been proposed as critical for Cas9’s rapid target site localization during interference. Additionally, it is plausible that the underwind feature of the CCR may facilitate PAM cleavage and subsequent recognition by the active center of Cas1 during spacer acquisition. Beyond the PAM-interacting domain, a cluster of basic residues on the REC2 domain of Cas9 (K221/R224/K226/K239/K255/K259/K260/K269) extensively contacts the DNA phosphate backbone (at bp +11 to +17), thereby enhancing DNA binding (Figure 3D). The PAM-distal region of the DNA further threads through the central channel of Csn2 tetramers, with its phosphate backbone (at bp +20 to +30) interacting with conserved basic residues K77, K131, and K160 (Figure 3D). These electrostatic interactions are non-specific, analogous to the interactions observed in DNA sliding clamps such as PCNA. Remarkably, the length of DNA from the beginning of CCR sequence to the last nucleotide protected by Csn2 is 30 bp, which precisely matches the length of prespacer required for integration by the Cas1-Cas2 complex, suggesting a potential ruler mechanism to determine the length of the prespacer at selection stage. Similar to the apo-state supercomplex, the association of Cas9 with Csn2 and Cas1 in the prespacer catching complex is exclusively mediated by its REC3 domain (residues 509-726) (Figure 3E). This domain interacts with both Cas1 and Csn2, presenting a buried surface area of approximately 1115 Å_2_. The REC3 domain is composed of twelve α-helices, with two distinct regions facilitating interactions with the Cas1 dimer. The loop connecting REC3’s α31 and α32 (residues Y679-S683) engages with conserved residues (E169, D171, K241) on α6 and α9 of Cas1_A_ (Figure 3F), while residues Y538, R542, E552 and Y576, 8 of REC3 extensively interact with the side chains of Q29, K17 and N18 of Cas1B (Figure 3G). Additionally, REC3 significantly contacts Csn2, with specific interactions between residues Q574, R577, N578, E579, and N581 of REC3 and N4, G40, F47, Q51 and K52 of the Csn2 head domain, creating a 520 Å_2_ interface (Figure 3G). Consequently, the Csn2 head domain is closely opposed to the N-terminal domain of Cas1(Figure 3H). Helix H11 of Cas1_B_ forms a π-π interaction and a hydrogen bond with F212 and D24 of Csn2, respectively (Figure 3I). Furthermore, Y213 of Csn2 establishes a hydrogen bond with the backbone carbonyl oxygen of S48 of Cas1_A_ (Figure 3J). Although these interaction interfaces are relatively small, they are crucial for complex stability; disruption of these interfaces results in a nearly complete loss of Cas9 association, indicating that the formation of this complex is primarily driven by protein-protein interactions rather than by protein-DNA interactions (Figure S7).

These findings align with prior observations that spacer acquisition preferentially selects prespacers with functional PAMs, dependent on Cas9’s PAM-interacting motif. By forming the prespacer catching complex with Csn2 and Cas1, Cas9 directs the complex to slide along the DNA, positioning Cas1 at an appropriate location relative to the PAM sequence.

### Cas9 dissociation reconfigures Csn2-Cas1-Cas2 to capture prespacer DNA

Within the same cryo-EM sample, we additionally identified a distinct particle population lacking Cas9, yielding a reconstruction at 3.76 Å resolution (Figure S5). This Cas9-dissociated subcomplex comprises a Csn2 tetramer bound to a Cas1 dimer, with a DNA duplex threaded through the central channel of the Csn2 tetramer and engaging a discrete density element (Figure 4A). The Cas1 dimer associated with Csn2 is well-resolved, enabling unambiguous structural modeling of the adjacent Cas2 C-terminal β-strand (Figure 4B and S8). Notably, the absence of density corresponding to the second Cas1 dimer and distal DNA regions suggests a highly dynamic or flexible conformation in this region.

**Figure 4.**
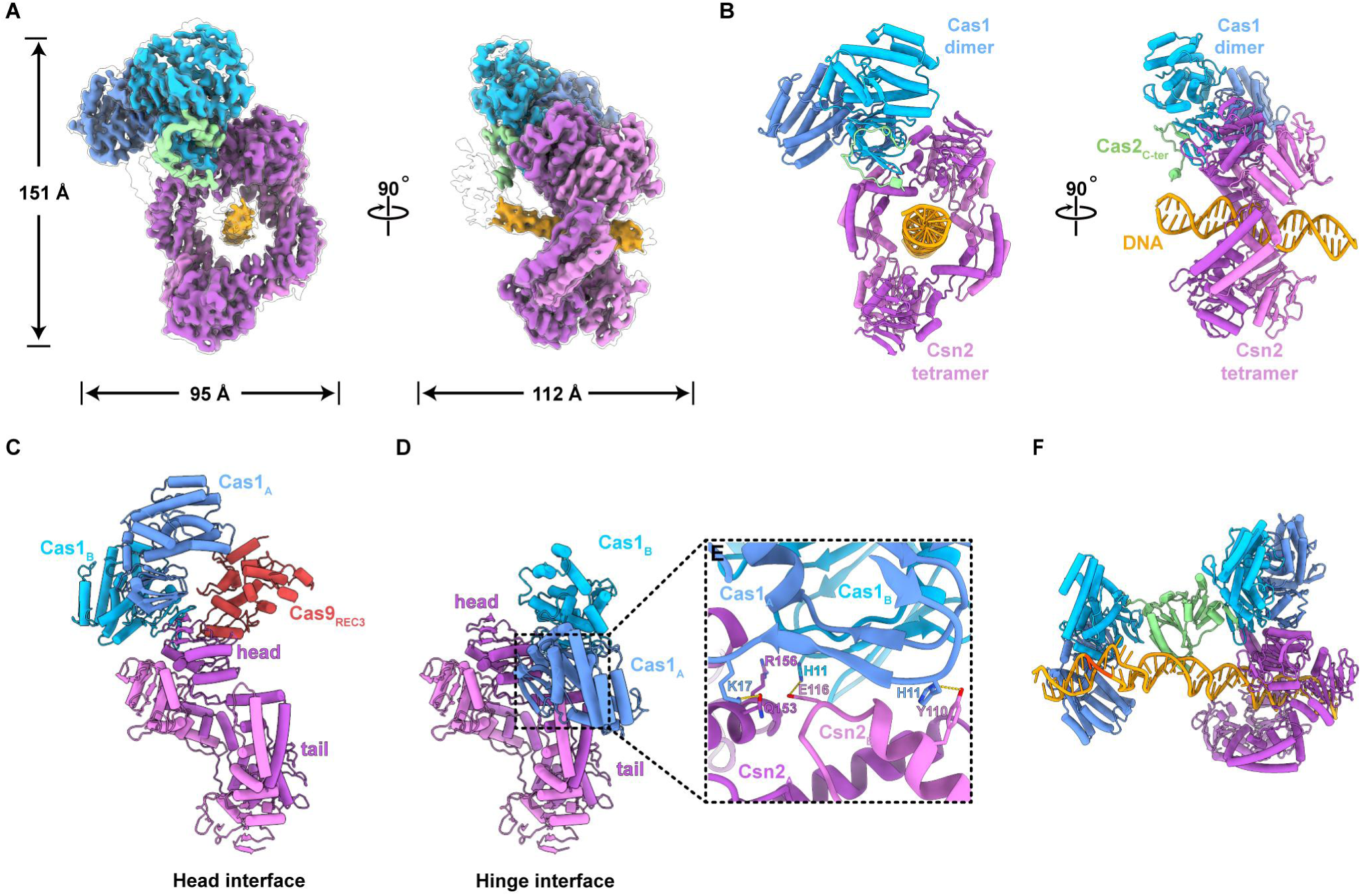
Cryo-EM structure of Cas9 dissociation subcomplex. **(A)** Cryo-EM density map of the Csn2-Cas1-Cas2 subcomplex. The raw electron density was shown as a transparent map. Subunits are colored as follows using sharpen map filtered to estimated overall resolution: Csn2, purple and violet; Cas1, blue and cyan; Cas2, green; DNA, orange. **(B)** The corresponding views of the molecular model. **(C)** The identical arrangement of the Csn2-Cas1 structure in the apo form and in the prespacer capture complex. **(D)** The distinct Csn2-Cas1 structure in the Cas9 dissociation subcomplex compared to the structure in **(C)**. **(E)** Close-up view of the interaction interface between Csn2 and Cas1 in the Cas9 dissociation subcomplex. **(F)** Alignment of Cas1-Cas2 hexamers with the Cas1 dimer in the Cas9-dissociated subcomplex positions Cas1 active sites near the defined PAM locus.

A defining feature of this Csn2-dsDNA-Cas1-Cas2 subcomplex, which distinguishes it from both the apo-state Cas9-Csn2-Cas1-Cas2 supercomplex and the prespacer-capturing complex, is the dissociation of Cas9. This structural rearrangement has two critical functional consequences. First, in the prespacer-capturing complex, Cas9 embeds the PAM-proximal DNA, rendering it inaccessible to Cas1, Cas2, or host nucleases. In contrast, the Cas9-dissociated subcomplex exposes the PAM-proximal prespacer DNA, potentially enabling host nucleases to process it and generate a 3′ overhang required for leader-side integration by Cas1. Second, in both the apo-state supercomplex and the prespacer-capturing complex, interactions between the Cas1 N-terminal domain and the Csn2 head domain are stabilized by Cas9’s REC3 domain. However, structural alignment of all Cas1-Cas2 hexamer structures with the Cas1 dimer in the prespacer-capturing complex revealed that Cas1-Cas2 hexamers cannot access DNA when Cas1 interacts with the Csn2 head domain (Figure S9). Cas9 dissociation disrupts the REC3/Cas1/Csn2 interface, abolishing the interaction between the Csn2 head domain and the Cas1 N-terminal domain (Figure 4C). This destabilization induces a sliding motion of the Cas1 dimer, leading to the formation of a new interface between the same Cas1 region and the hinge connecting the Csn2 head and tail domains—a configuration also observed in the StCsn2-dsDNA-Cas1-Cas2 dimer (PDB: 6QXT) (Figure 4D and Video S1). At this interface, Y110 and E116 of Csn2_B_ forms π-π stacking and hydrogen bond with H11 of Cas1_A_ and Cas1_B_, respectively. Additional hydrogen bond is also observed between Q153 of Csn2_A_ and K17 of Cas1_A_ (Figure 4E). R156 of Csn2_A_ also forms π-π stacking with H11 of Cas1_B_ (Figure 4E). Although this interface is modest in size (1,248 Å_2_), direct DNA binding by Cas2 stabilizes the association of Cas1-Cas2 hexamers. Importantly, alignment of Cas1-Cas2 hexamer structures with the Cas1 dimer in the Cas9-dissociated subcomplex positions the Cas1 active sites in close proximity to the previously defined PAM locus. The structural variability introduced by the flexible C-terminal loop of Cas2 may further enable the Cas1 dimer to engage the processed 3′ overhang following PAM cleavage, facilitating leader-side integration (Figure 4F).

Together, Cas9 dissociation triggers a structural reconfiguration of the Csn2-Cas1-Cas2 assembly, relocating Cas1 to the Csn2 hinge region. This rearrangement exposes the PAM-proximal DNA, permitting Cas1-Cas2 hexamers to access the DNA and coordinate prespacer processing and integration.

## Discussion

Based on our structural findings, we propose a model for spacer acquisition in type II-A CRISPR-Cas systems (Figure 5). The Cas9_2_-Csn2_8_-Cas1_8_-Cas2_4_ supercomplex represents a resting state, designed to minimize off-target cleavage and unintended integration while maintaining the ability to survey free DNA ends. Upon encountering free DNA ends that generated from phage or plasmid DNA, Csn2 initiates the process by encircling and threading the DNA end into the supercomplex. DNA binding induces a conformational change in Cas9, triggering the dissociation of the supercomplex and the formation of the prespacer catching complex. This complex then slides along the DNA until it encounters a PAM sequence. At this juncture, Cas9 is released, exposing the PAM-proximal prespacer DNA. Subsequently, an unidentified host nuclease processes this region, cleaving the PAM sequence and generating a 3’ overhang. Concurrently, the Cas1 dimer translocate to the hinge connecting the Csn2 head and tail domains, allowing Cas1-Cas2 to engage the 3’ overhang for leader-side integration. A second, yet-to-be-identified host nuclease then cleaves at the PAM-distal end, which is encapsulated by Csn2, leading to Csn2 dissociation and completing spacer-side integration.

**Figure 5.**
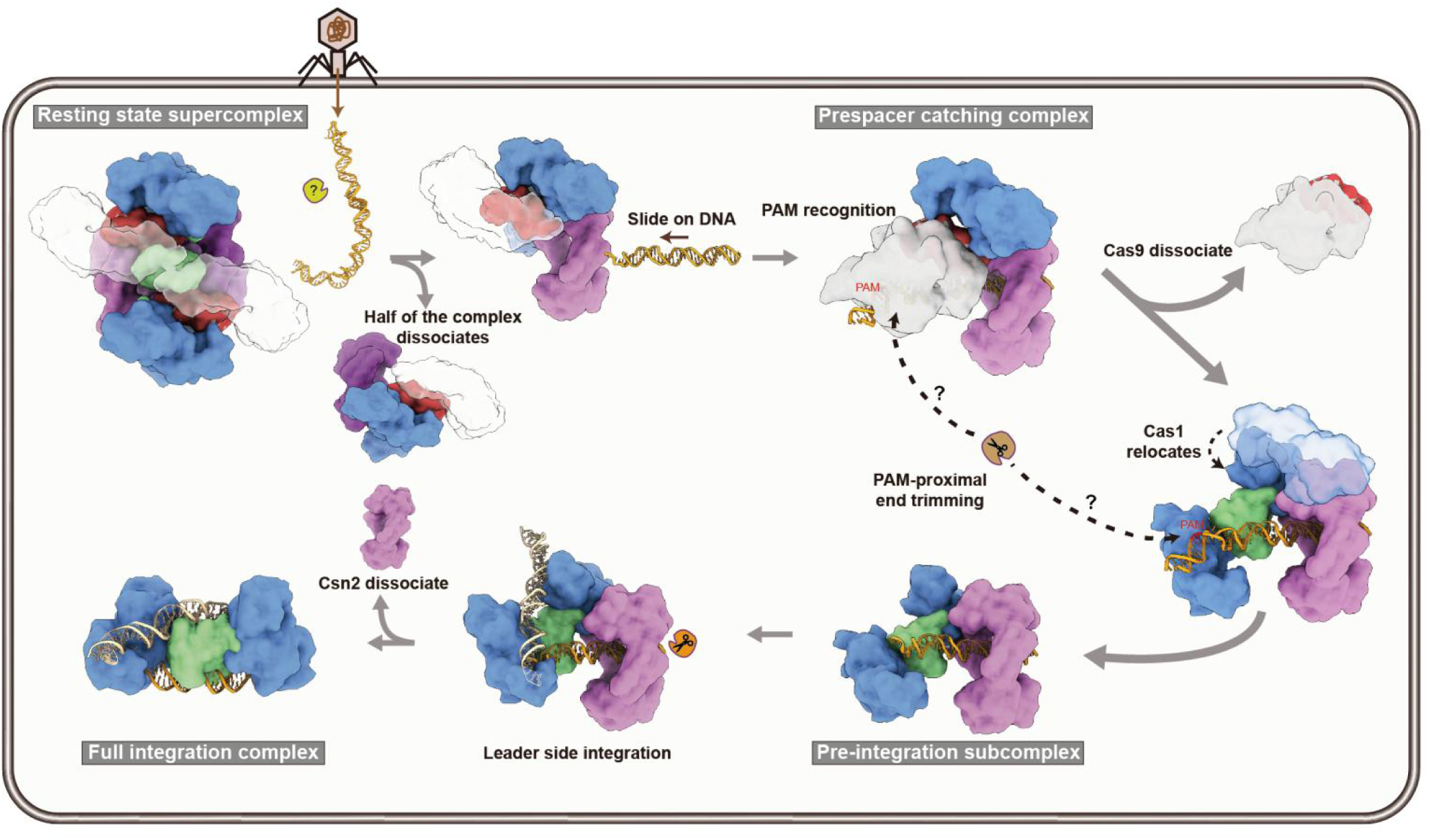
Model of spacer acquisition in type II-A CRISPR-Cas system. In the resting state, the Cas9-Csn2-Cas1-Cas4 supercomplex minimizes off-target cleavage while surveying free DNA ends. Upon encountering free DNA, Csn2 threads the DNA into the supercomplex, inducing a conformational change in Cas9 that triggers dissociation and forms the prespacer-capturing complex. The complex slides along the DNA until it reaches a PAM sequence, releasing Cas9 and exposing the PAM-proximal prespacer. This region is processed by a host nuclease, generating a 3′ overhang. Cas1-Cas2 then engages the overhang for leader-side integration, and another host nuclease cleaves the PAM-distal end, leading to Csn2 dissociation and spacer-side integration.

Our findings provide a comprehensive structural framework for understanding Cas9 directed prespacer selection in type II-A CRISPR-Cas systems. The dynamic interplay between Cas9, Csn2, and Cas1-Cas2 ensures precise and efficient spacer acquisition. The observed structural transitions highlight the importance of allosteric regulation and protein-protein interactions in coordinating the complex steps of CRISPR adaptive immunity. Recent studies in type I systems have elucidated Cas1 or Cas4-dependent PAM recognition and sequential cleavage mechanisms, ensuring directional integration with the prespacer’s PAM-distal 3’ end being trimmed and integrated first.^13–23^ In contrast, the type II-A system presents an inverse scenario, where the PAM-proximal end must be trimmed and integrated first. While *Spy*Cas9 has demonstrated prespacer trimming activity, this function is dispensable in vivo.^40^ Therefore, identifying the host nucleases responsible for PAM trimming is crucial for a complete understanding of the interplay between prespacer processing and integration orientation in type II-A adaptation.

## Acknowledgements

We thank the Instrument Analysis Center (IAC) at Shanghai Jiao Tong University for cryo-EM data collection. This work was supported by National Key Research and Development Program of China (grant 2023YFC3402300 to M.L.), STI2030-Major Projects (2021ZD0203400,Y.X.), National Natural Science Foundation of China (82473977, Y.X.; 32271330, M.C.; 82304614, Z.L.), Natural Science Foundation of Jiangsu Province (SBK2024010634, Y.X.), the Project Program of State Key Laboratory of Natural Medicines, China Pharmaceutical University (SKLNMZZ2024JS31, Y.X.), the National University of Singapore Start-up grant (M.L.), the Ministry of Education Tier 2 Grant (MOE-T2EP30222-001, M.L.) and the National Research Foundation grant (NRF-CRP22-2019-0001, M.L.).

## Author contributions

Z.X.L., Y.T.L., M.L.L., M.L. and Y.B.X. conceived the project and designed the experiments. Z.X.L., Y.T.L., J.P.K., Q.Q.W., P.P.H., Y.F.L., H.F.L., L.Q.H., G.Y.L., T.Z, C.Y.H., Y.T.H and Y.Z. carried out the experiments. Z.X.L., Y.T.L., M.R.C., C.Y.H., Z.H.Y., M.L.L., M.L. and Y.B.X. analyzed the data. Z.X.L., M.L.L., M.L. and Y.B.X. wrote the manuscript. All authors discussed the results and contributed to the final manuscript.

## Additional information

**Supplementary information** is available for this paper.

**Correspondence and requests for materials** should be addressed to Xiao, Y.

**Peer review information**

## RESOURCE AVAILABILITY

### Lead contact

Further information and requests for resources and reagents should be directed to and will be fulfilled by the lead contact, Yibei, Xiao.

### Materials availability

This study did not generate new unique reagents.

### Data availability

The atomic coordinates of the reported cryo-EM structures have been deposited in the Protein Data Bank under accession numbers 92JD, 9J2S, 9J2I and 9J2I, respectively. The corresponding cryo-EM density maps have been deposited in the Electron Microscopy Data Bank under accession codes EMD-34441, EMD-61100, EMD-61101 and EMD-61097, respectively.

These data are publicly available as of the date of publication. This paper does not report original code.

Any additional information required to reanalyze the data reported in this paper is available from the lead contact upon request.

## EXPERIMENTAL MODEL AND SUBJECT DETAILS

### Microbes

*Escherichia coli* cells were cultured in LB medium.

## METHODS DETAILS

### Cloning, expression, and purification

Full-length *cas9*, *csn2*, *cas1,* and *cas2* cDNAs were PCR amplified from *E. faecalis* genomic DNA. The *cas9* gene was cloned into pET24a vector with a C-terminal His_6_ tag. The *csn2* gene was cloned into a pSJ8 vector containing an N-terminal His_6_-MBP tag and a TEV protease cleavage site. Plasmids encoding Cas1 and Cas2 were constructed as described previously^8^. Sequence-verified plasmids were transformed into *E. coli* BL21 Star (DE3) cells. The transformed cells were grown at 37°C in LB medium until the optical density at 600 nm (OD_600_) reached 0.6∼0.8. Protein expression was induced by adding Isopropyl β-D-1-Thiogalactopyranoside (IPTG) to a final concentration of 0.5 mM at 25°C overnight. Cells were harvested by centrifugation and lysed using a high-pressure homogenizer in a buffer containing 50 mM HEPES (pH 7.5), 20 mM imidazole, and 500 mM NaCl. The lysate was centrifuged twice at 15,000 r.p.m. for 30 min at 4°C, and the supernatant was applied onto a pre-equilibrated Ni-NTA column (Smart-Lifesciences). The column was washed with 200 mL buffer containing 20 mM HEPES (pH 7.5) and 500 mM NaCl, and bound protein was eluted using a buffer containing 20 mM HEPES (pH 7.5), 500 mM NaCl, and 300 mM imidazole. The His_6_-MBP tag on Csn2 was cleaved using TEV protease at 4°C overnight. Cas9 and tag-cleaved Csn2 were further purified using a HiLoad 16/600 Superdex 200 column (Cytiva) equilibrated in 20 mM HEPES (pH 7.5) and 250 mM NaCl. The His_6_-Twin-Strep-SUMO tag on Cas1 and Cas2 was cleaved using SUMO protease at 4°C overnight. The tag-cleaved Cas1 and Cas2 proteins were then mixed at a 2:1 molar ratio and further purified using a HiLoad 16/600 Superdex 200 column (Cytiva) equilibrated in 20 mM HEPES (pH 7.5) and 500 mM NaCl. The corresponding peak fractions were pooled and snap-frozen in liquid nitrogen for further use. All mutants were constructed using rolling circle replication PCR, and their purification followed the same protocol as that of the wild type protein.

### Nucleic acid preparation

Plasmid DNA for in vitro cleavage assays was generated by cloning annealed complementary oligonucleotides into the pETDuet-1 vector. The resulting plasmid was digested with the ScaI restriction enzyme to produce linearized target DNA. The RNAs used in this study were synthesized by GenScript. The DNA oligonucleotides were synthesized by Sangon Biotech. DNA duplexes were annealed by heating at 95 °C, followed by slow cooling to room temperature in an annealing buffer containing 20 mM HEPES (pH 7.5) and 250 mM NaCl.

### In vitro DNA cleavage assays

Cas9 and a 1.5-fold molar excess of tracrRNA/crRNA were incubated at 37 °C for 5 min to assemble the Cas9-RNA complex. Cleavage reactions were initialized by adding 50 nM Cas9-RNA and 10 nM circular or linearized target DNA in a reaction buffer containing 20 mM HEPES (pH 7.5), 250 mM NaCl, and 10 mM MgCl_2_. After incubation at 37°C for 30 min, the reactions were quenched by adding a 5× loading solution containing 30% glycerol, 1.2% SDS, and 250 mM EDTA. Reaction products were analyzed by 1% agarose gel electrophoresis, and bands were visualized by pre-staining with GelRed. To investigate the effect of the Cas1-Cas2 complex and Csn2 on the cleavage activity of Cas9-RNA, 400 nM Csn2 tetramer, 400 nM Cas1-Cas2 hexamer, or both were first incubated with 50 nM Cas9-RNA at room temperature for 30 min. Subsequent steps followed the same protocol as described above.

### Gel filtration assay

Analytical size exclusion chromatography (SEC) was performed using a Superdex 200 Increase 10/300 GL column (Cytiva). The column was equilibrated with buffer containing 20 mM HEPES (pH 7.5) and 250 mM NaCl. To assess complex formation, protein samples were mixed and gradually exchanged into buffer containing 20 mM HEPES (pH 7.5) and 250 mM NaCl before loading onto the column. Elution fractions were analyzed by 8∼20% sodium dodecyl sulfate polyacrylamide gel electrophoresis (SDS-PAGE) and visualized by Coomassie staining.

### Electron microscopy sample preparation and data collection

To assemble the *E. faecalis* Cas9/RNA -Csn2-Cas1-Cas2 supercomplex, Cas9 was first incubated with a 1.5-fold molar excess of tracrRNA/crRNA at 37 °C for 5 min. Subsequently, appropriate amounts of Csn2, Cas1, and Cas2 were added, and the mixture was incubated for an additional 30 min. The resulting supercomplex was purified by SEC using a Superdex 200 Increase 10/300 GL column (Cytiva) in buffer containing 20 mM HEPES (pH 7.5) and 250 mM NaCl. For cryo-EM grid preparation of the Cas9/RNA-Csn2-Cas1-Cas2 supercomplex, a 3.5 μL aliquot of the purified supercomplex (0.6 mg/mL) was applied to glow-discharged Quantifoil holey gold grids (R 1.2/1.3, Au 300 mesh), followed by an incubation for 15 s. Grids were blotted for 1 s under 100% relative humidity and plunge-frozen in liquid ethane cooled by liquid nitrogen using a Vitrobot System. A total of 617 movies were collected using a FEI Titan Krios G3i (Thermo Fisher) operating at 300 kV and equipped with a K2 Summit direct electron detector. All cryo-EM movies were recorded in counting mode using SerialEM.^46^

To assemble the *E. faecalis* Cas9/RNA-Csn2-Cas1-Cas2-DNA complex, the DNA substrate was incubated with the preassembled Cas9/RNA-Csn2-Cas1-Cas2 supercomplex at a 1.5:1 molar ratio (DNA: Cas9) at 4 °C for 30 min. An aliquot of 3.5 μL of the sample was then applied to glow-discharged Quantifoil holey gold grids (R 1.2/1.3, Au 300 mesh), followed by a 15 s waiting time and a 3-5 s blotting time with the blot force set to 2. The grids were plunge-frozen in liquid ethane using a Virtrobot Mark IV (Thermo Fisher). Movies were collected on an FEI Titan Krios G3i (Thermo Fisher) operated at 300 kV and equipped with a Falcon 4i detector. For this dataset, movies were recorded in counting mode with GPU acceleration. Detailed parameters, including electron dose, pixel size, and magnifications settings, are provided in Supplementary Data Table 1.

### EM data processing

Images were processed using cryoSPARC (v.4.2) unless otherwise indicated.^47^ Movie frames were aligned and summed using MotionCor2.^48^ Contrast transfer function (CTF) parameters were estimated for individual particles on each micrograph using cryoSPARC. After patch CTF estimation, micrographs with a resolution estimate worse than 6 Å were discarded. Blob picker was used for particle picking, and the selected particles were inspected manually. Particles with NCC scores below 0.3 were discarded. 2D classification, 3D classification, and 3D refinement were also performed in cryoSPARC. All refinements followed the gold-standard procedure, in which two half-datasets were refined independently. The overall resolutions were estimated based on the gold-standard Fourier shell correlation (FSC) criterion of 0.143. Local resolutions were calculated in cryoSPARC and visualized using ChimeraX.^49^

For Dataset 1 [Cas9/RNA-Csn2-Cas1-Cas2, apo state], 617 movies were recorded, and 611 images were retained after manual inspection. Following 2D classification, 145,992 particles were selected from 542,743 auto-picked particles and subjected to 3D reconstruction and classification. To improve resolution, 89,654 particles belonging to the two best 3D classes were retained for an additional round of 3D refinement, from which 69,551 particles in the best class were selected. Further 3D refinement, followed by non-uniform refinement, local CTF refinement, duplicate removal, and post-processing, resulted in a final 3D map with an average resolution of 3.90 Å (EMD-34441).

For Dataset 2 [Cas9/RNA-Csn2-Cas1-Cas2 with DNA substrate], 15,941 movies were recorded, and 15,719 micrographs were retained after manual curation. From 4,217,921 auto-picked particles, 1,056,302 particles were selected following 2D classification for subsequent 3D reconstruction and classification. Two distinct stable complexes were identified and selected for further refinement. Iterative 3D refinement combined with non-uniform refinement and post-processing yielded final 3D maps with average resolutions of 2.96 Å (EMD-61100), 3.01 Å (EMD-61101) and 3.76 Å (EMD-61097), respectively.

Detailed data processing and refinement statistics for the cryo-EM structures are summarized in Extended Data Table 1. Reconstructions were sharpened and filter to overall resolution by using DeepEMhancer with the default tight mask pre-set.^50^

### Model building and refinement

Crystal structures of *Efa*Csn2(PDB 3S5U) and *Efa*Cas1-Cas2(PDB 5XVN) were used for docking in this study. An initial model of *Efa*Cas9 was generated using AlphaFold3.^51^ This model was initially docked into the cryo-EM density map using UCSF Chimera X^49^ and manually rebuilt on Coot.^52^ Regions exhibiting poorly resolved density, only fitting in map was performed without subsequent manual refinement. The models were further adjusted in Coot and refined using phenix.real_space_refine.^49, 53^ Model quality was assessed using MolProbity in Phenix.^54^

## References

1 Barrangou, R. et al. CRISPR provides acquired resistance against viruses in prokaryotes. Science 315, 1709–1712 (2007).

2 Marraffini, L. A. & Sontheimer, E. J. CRISPR interference: RNA-directed adaptive immunity in bacteria and archaea. Nat. Rev. Genet. 11, 181–190 (2010).

3 Marraffini, L. A. CRISPR-Cas immunity in prokaryotes. Nature 526, 55–61 (2015).

4 Jackson, S. A. Et al. CRISPR-Cas: Adapting to change. Science 356, eaal5056 (2017).

5 Pacesa, M., Pelea, O. & Jinek, M. Past, present, and future of CRISPR genome editing technologies. Cell 187, 1076–1100 (2024).

6 McGinn, J. & Marraffini, L. A. Molecular mechanisms of CRISPR-Cas spacer acquisition. Nat. Rev. Microbiol. 17, 7–12 (2019).

7 Wang, J. et al. Structural and Mechanistic Basis of PAM-dependent spacer acquisition in CRISPR-Cas Systems. Cell 163, 840–853 (2015).

8 Xiao, Y., Ng, S., Nam, K. H. & Ke, A. How type II CRISPR-Cas establish immunity through Cas1-Cas2 mediated spacer integration. Nature 550, 137–141 (2017).

9 Wright, A. V. Et al. Structures of the CRISPR genome integration complex. Science 357, 1113–1118 (2017).

10 Yosef, I., Goren, M. G. & Qimron, U. Proteins and DNA elements essential for the CRISPR adaptation process in *Escherichia coli*. Nucleic Acids Res. 40, 5569–5576 (2012).

11 Nuñez, J. K. Et al. Cas1-Cas2 complex formation mediates spacer acquisition during CRISPR-Cas adaptive immunity. Nat. Struct. Mol. Biol. 21, 528–534 (2014).

12 Nuñez, J. K., Lee, A. S. Y., Engelman, A. & Doudna, J. A. Integrase-mediated spacer acquisition during CRISPR-Cas adaptive immunity. Nature 519, 193–198 (2015).

13 Kieper, S. N. Et al. Cas4 facilitates PAM-compatible spacer selection during CRISPR adaptation. Cell Rep. 22, 3377–3384 (2018).

14 Lee, H., Zhou, Y., Taylor, D. W. & Sashital, D. G. Cas4-dependent prespacer processing ensures high-fidelity programming of CRISPR arrays. Mol. Cell 70, 48–59 (2018).

15 Rollie, C., Graham, S., Rouillon, C. & White, M. F. Prespacer processing and specific integration in a type I-A CRISPR system. Nucleic Acids Res. 46, 1007–1020 (2017).

16 Masami, S., Garrett, S. C., Graveley, B. R. & Terns, M. P. Cas4 nucleases define the PAM, length, and orientation of DNA fragments integrated at CRISPR loci. Mol. Cell 70, 814–824.e816 (2018).

17 Kieper, S. N. et al. Cas4-Cas1 is a Protospacer adjacent motif-processing factor mediating half-site spacer integration during CRISPR adaptation. CRISPR J. 4, 536–548 (2021).

18 Hu, C. et al. Mechanism for Cas4-assisted directional spacer acquisition in CRISPR-Cas. Nature 598, 515–520 (2021).

19 Dhingra, Y., Suresh, S. K., Juneja, P. & Sashital, D. G. PAM binding ensures orientational integration during Cas4-Cas1-Cas2-mediated CRISPR adaptation. Mol. Cell 82, 4353–4367 (2022).

20 Almendros, C., Nobrega, F. L., McKenzie, R. E. & Brouns, S. J. J. Cas4-Cas1 fusions drive efficient PAM selection and control CRISPR adaptation. Nucleic Acids Res. 47, 5223–5230 (2019).

21 Kim, S. et al. Selective loading and processing of prespacers for precise CRISPR adaptation. Nature 579, 141–145 (2020).

22 Shiriaeva, A. A. Et al. Host nucleases generate prespacers for primed adaptation in the *E. coli* type I-E CRISPR-Cas system. Sci. Adv. 8, eabn8650 (2022).

23 Drabavicius, G. et al. DnaQ exonuclease-like domain of Cas2 promotes spacer integration in a type I-E CRISPR-Cas system. EMBO Rep. 19, e45543 (2018).

24 Mojica, F. J. M., Díez-Villaseñor, C., García-Martínez, J. & Almendros, C. Short motif sequences determine the targets of the prokaryotic CRISPR defence system. Microbiology (Reading) 155, 733–740 (2009).

25 Marraffini, L. A. & Sontheimer, E. J. Self versus non-self discrimination during CRISPR RNA-directed immunity. Nature 463, 568–571 (2010).

26 Lee, H., Dhingra, Y. &, D. G. The Cas4-Cas1-Cas2 complex mediates precise prespacer processing during CRISPR adaptation. Elife 8, e44248 (2019).

27 Zhang, J., Kasciukovic, T. & White, M. F. The CRISPR associated protein Cas4 is a 5’ to 3’ DNA exonuclease with an iron-sulfur cluster. PloS One. 7, e47232 (2012).

28 Cristóbal, A., Nobrega, F. L., Mckenzie, R. E. & Brouns, S. Cas4-Cas1 fusions drive efficient PAM selection and control CRISPR adaptation. Nucleic Acids Res. 47, 5223–5230 (2019).

29 Heler, R. et al. Cas9 specifies functional viral targets during CRISPR-Cas adaptation. Nature 519, 199–202 (2015).

30 Wei, Y., Terns, R. M. & Terns, M. P. Cas9 function and host genome sampling in Type II-A CRISPR-Cas adaptation. Genes Dev. 29, 356–361 (2015).

31 Wilkinson, M. et al. Structure of the DNA-bound spacer capture complex of a type II CRISPR-Cas system. Mol. Cell 75, 90–101 (2019).

32 Arslan, Z. et al. Double-strand DNA end-binding and sliding of the toroidal CRISPR-associated protein Csn2. Nucleic Acids Res. 41, 6347–6359 (2013).

33 Ellinger, P. et al. The crystal structure of the CRISPR-associated protein Csn2 from *Streptococcus agalactiae*. J. Struct. Biol. 178, 350–362 (2012).

34 Lee, K. H. et al. Identification, structural, and biochemical characterization of a group of large Csn2 proteins involved in CRISPR-mediated bacterial immunity. Proteins. 80, 2573–2582 (2012).

35 Nam, K. H., Kurinov, I. & Ke, A. Crystal structure of clustered regularly interspaced short palindromic repeats (CRISPR)-associated Csn2 protein revealed Ca^2+^-dependent double-stranded DNA binding activity. J. Biol. Chem. 286, 30759–30768 (2011).

36 Wright, A. V. & Doudna, J. A. Protecting genome integrity during CRISPR immune adaptation. Nat. Struct. Mol. Biol. 23, 876–883 (2016).

37 Budhathoki, J. B. Et al. Real-time observation of CRISPR spacer acquisition by Cas1-Cas2 integrase. Nat. Struct. Mol. Biol. 27, 489–499 (2020).

38 Ka, D. et al. Crystal structure of *Streptococcus pyogenes* Cas1 and its interaction with Csn2 in the type II CRISPR-Cas system. Structure 24, 70–79 (2016).

39 Ka, D., Jang, D. M., Han, B. W. & Bae, E. Molecular organization of the type II-A CRISPR adaptation module and its interaction with Cas9 via Csn2. Nucleic Acids Res. 46, 9805–9815 (2018).

40 Jakhanwal, S. et al. A CRISPR-Cas9-integrase complex generates precise DNA fragments for genome integration. Nucleic Acids Res. 49, 3546–3556 (2021).

41 Zuo, Z. & Liu, Jin. Allosteric Regulation of CRISPR-Cas9 for DNA Targeting and Cleavage. Curr. Opin. Struct. Biol. 62, 166–174 (2020).

42 Chen, J.S. et al. Enhanced proofreading governs CRISPR-Cas9 targeting accuracy. Nature 550, 407–410 (2017).

43 Zhu, X. et al. Cryo-EM structures reveal coordinated domain motions that govern DNA cleavage by Cas9. Nat. Struct. Mol. Biol. 26, 679–685 (2019).

44 Chen, J. S. Et al. Enhanced proofreading governs CRISPR-Cas9 targeting accuracy. Nature 550, 407–410 (2017).

45 Shams, A. et al. Comprehensive deletion landscape of CRISPR-Cas9 identifies minimal RNA-guided DNA-binding modules. Nat. Commun. 12, 5664 (2021).

46 Mastronarde, D. N. Automated electron microscope tomography using robust prediction of specimen movements. J. Struct. Biol. 152, 36–51 (2005).

47 Punjani, A., Rubinstein, J. L., Fleet, D. J. & Brubaker, M. A. cryoSPARC: algorithms for rapid unsupervised cryo-EM structure determination. Nat. Methods 14, 290–296 (2017).

48 Zheng, S. Q. Et al. MotionCor2: anisotropic correction of beam-induced motion for improved cryo-electron microscopy. Nat. Methods 14, 331–332 (2017).

49 Goddard, T. D. Et al. UCSF ChimeraX: Meeting modern challenges in visualization and analysis. Protein Sci. 27, 14–25 (2018).

50 Sanchez-Garcia, R., Gomez-Blanco, J., Cuervo, A., Carazo, J. M., Sorzano, C. O. S., & Vargas, J. (2021). DeepEMhancer: a deep learning solution for cryo-EM volume post-processing. Communications biology 4(1), 874.

51 Abramson, J. et al. (2024). Accurate structure prediction of biomolecular interactions with AlphaFold 3. Nature 630, 493 – 500. 10.1038/s41586-024-07487-w.

52 Emsley, P., Lohkamp, B., Scott, W. G. & Cowtan, K. Features and development of Coot. Acta Crystallogr. D. Biol. Crystallogr. 66, 486–501 (2010).

53 Afonine, P. V. Et al. Real-space refinement in PHENIX for cryo-EM and crystallography. Acta Crystallogr. D. Struct. Biol. 74, 531–544 (2018).

54 Williams, C. J. Et al. MolProbity: More and better reference data for improved all-atom structure validation. Protein Sci. 27, 293–315 (2018).

