## Supplemental figure and table for "Structural Basis for Cas9-Directed Spacer Acquisition in Type II-A CRISPR-Cas Systems"

A

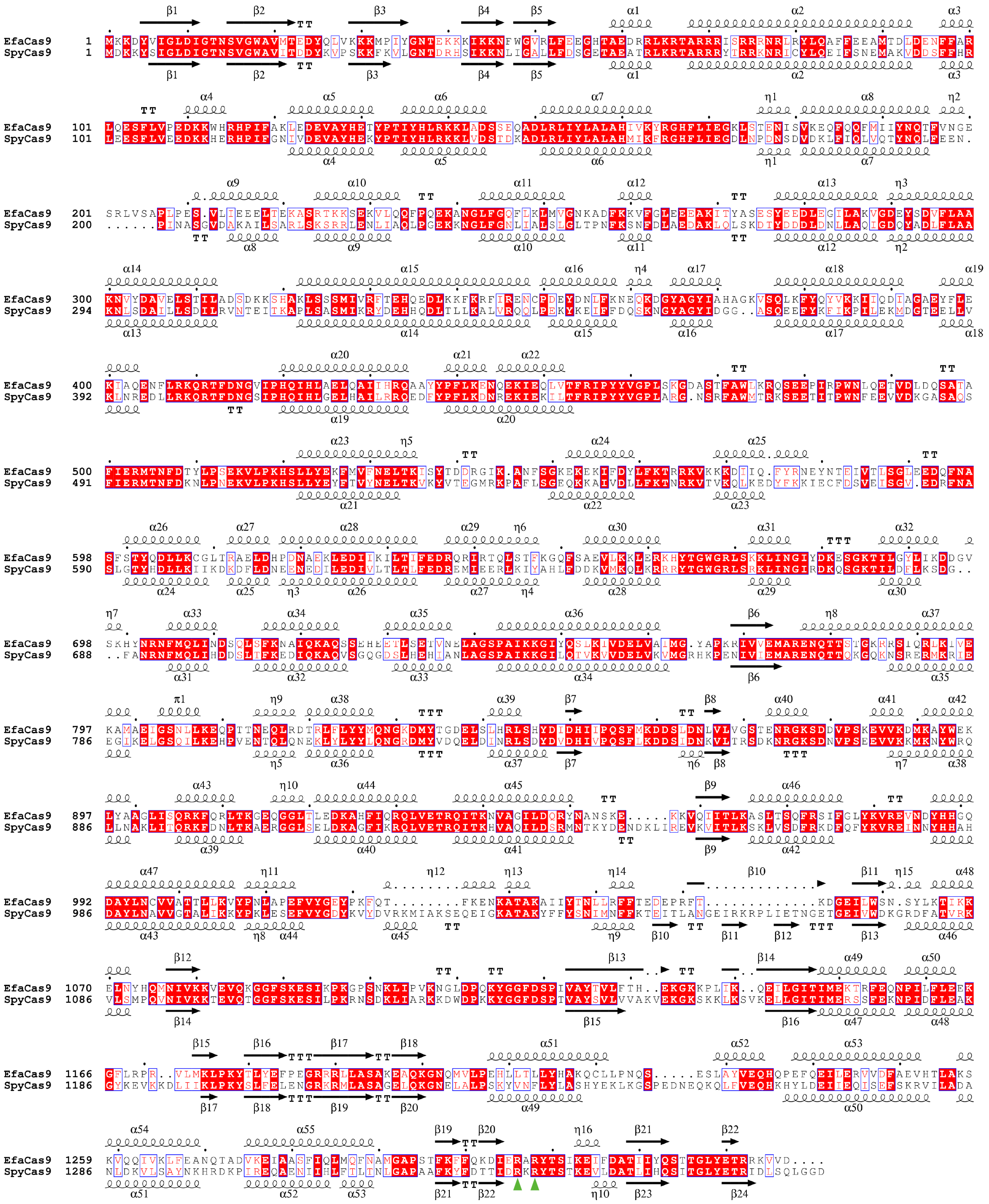
EfaCas9
SpyCas9

B

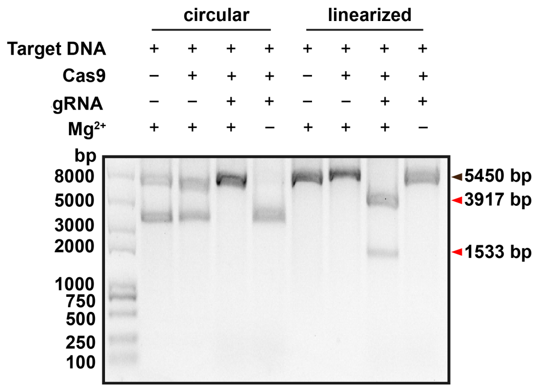
circular
linearized
Target DNA
Cas9
gRNA
Mg²⁺
bp
8000
5000
3000
2000
1000
750
500
250
100
5450 bp
3917 bp
1533 bp

**Figure S1 Sequence alignment and nuclease activity of EfaCas9 compared to SpyCas9, related to Figure 1**

**(A)** Sequence alignment of EfaCas9 and SpyCas9. Homologs from Enterococcus faecalis TX0027 (accession code: A0A6I4XW76) and Streptococcus pyogenes serotype M1 (accession code: Q99ZW2) are included in the alignment. Identical residues are boxed in red. The two residues responsible for PAM recognition are marked with green triangles.

**(B)** EfaCas9/gRNA cleaves both circular and linearized plasmid DNA containing a complementary target sequence and a 5’-NGG-3’ PAM.

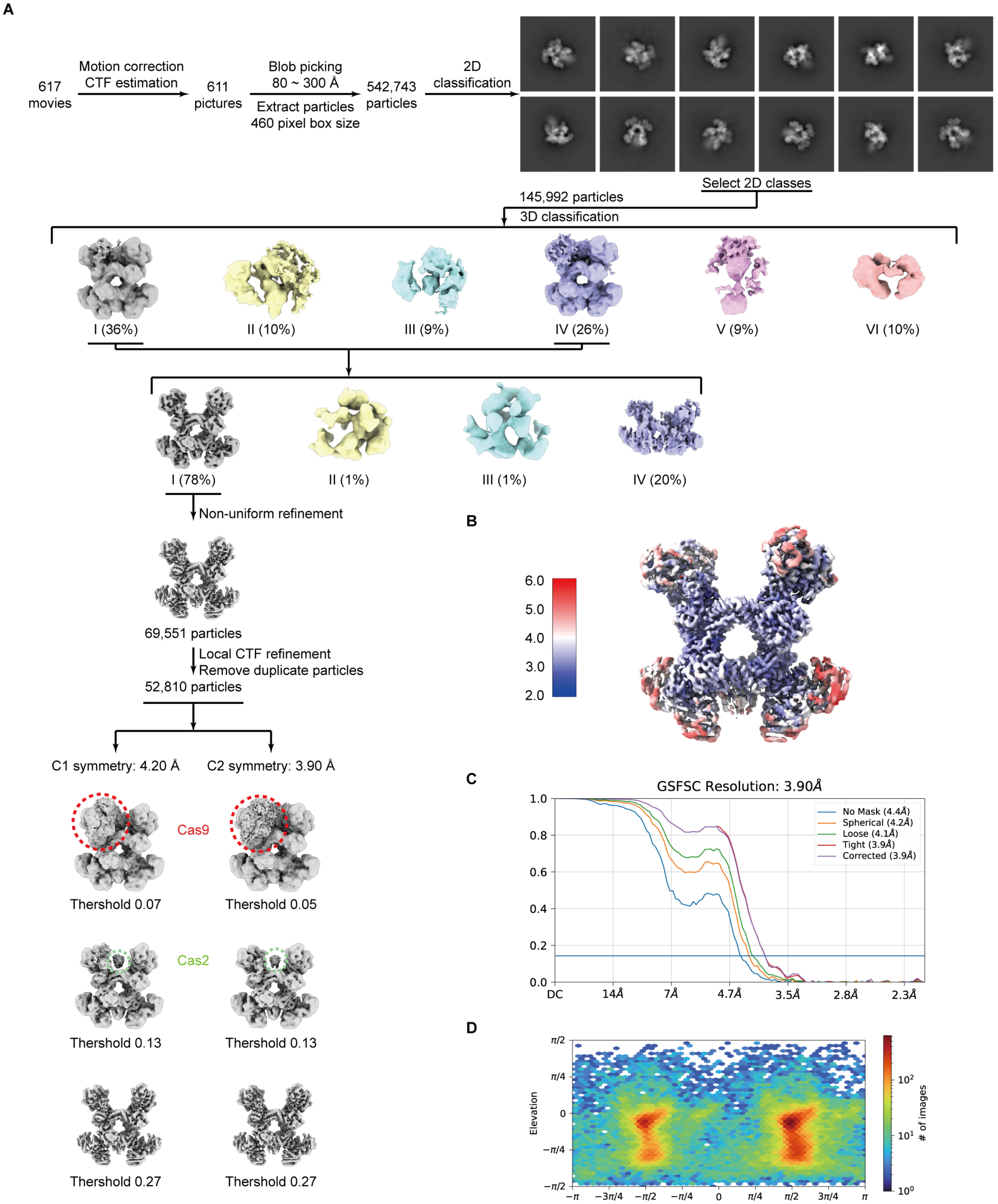
A
617 movies
Motion correction
CTF estimation
611 pictures
Blob picking
80 ~ 300 Å
Extract particles
460 pixel box size
542,743 particles
2D classification
Select 2D classes
145,992 particles
3D classification
I (36%)
II (10%)
III (9%)
IV (26%)
V (9%)
VI (10%)
I (78%)
II (1%)
III (1%)
IV (20%)
Non-uniform refinement
69,551 particles
Local CTF refinement
Remove duplicate particles
52,810 particles
C1 symmetry: 4.20 Å
C2 symmetry: 3.90 Å
Cas9
Thershold 0.07
Thershold 0.05
Cas2
Thershold 0.13
Thershold 0.13
Thershold 0.27
Thershold 0.27
B
6.0
5.0
4.0
3.0
2.0
C
GSFSC Resolution: 3.90Å
No Mask (4.4Å)
Spherical (4.2Å)
Loose (4.1Å)
Tight (3.9Å)
Corrected (3.9Å)
1.0
0.8
0.6
0.4
0.2
0.0
DC
14Å
7Å
4.7Å
3.5Å
2.8Å
2.3Å
D
Elevation
Azimuth
### of images

**Figure S2 Single-particle cryo-EM analysis of the apo Cas9/RNA-Csn2-Cas1-Cas2 supercomplex.**

**(A)** Cryo-EM reconstruction workflow for the apo Cas9/RNA-Csn2-Cas1-Cas2 supercomplex.

**(B-D)** Cryo-EM map of the Cas9/RNA-Csn2-Cas1-Cas2 supercomplex filtered to their estimated overall resolution and colored according to local resolution (**B**). The FSC curves (**C**) and the viewing direction distribution plot (**D**) for data processing.

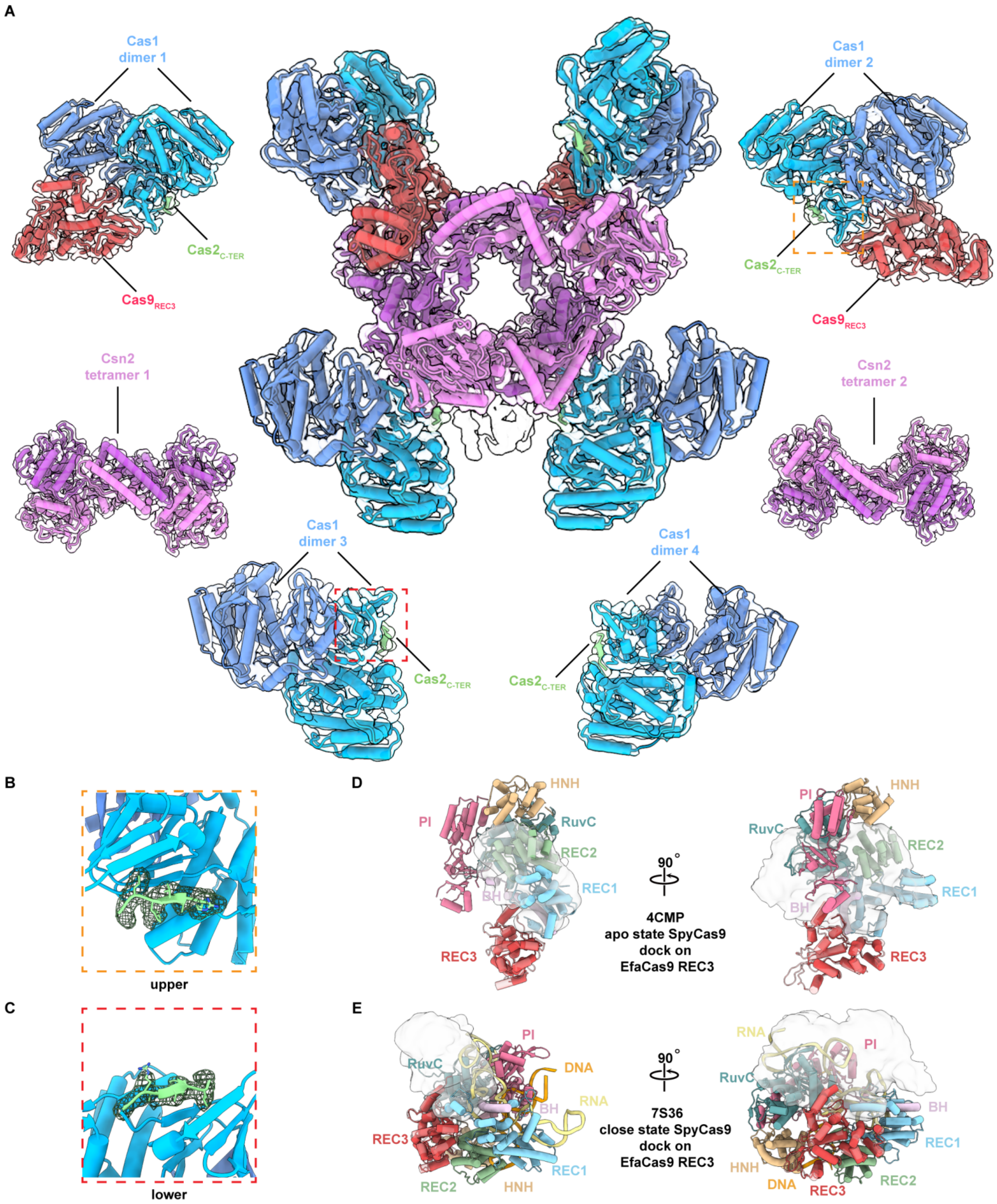


**Figure S3 The cryo-EM structure of apo Cas9/RNA-Csn2-Cas1-Cas2 supercomplex.**

**(A)** The structure model is shown in cartoon with sharpened cryo-EM map shown as a transparent surface. The supercomplex structure modeled here contains all four proteins with the stoichiometry $Cas9_2$-$Csn2_8$-$Cas1_8$-$Cas2_4$ . The REC3 domain (red) of the two Cas9 monomers has been modeled as well. Within each of the four Cas2 (green), only one β-fold that directly interact with Cas1 can be built.

**(B-C)** The β-sheet region of Cas2 adjacent to Cas1 can be clearly modeled with well-defined density.

**(D-E)** The density belonging to Cas9 could not be aligned to existing SpyCas9 structures, apo-state (PDB: 4CMP) or closed-state (PDB:7S36).

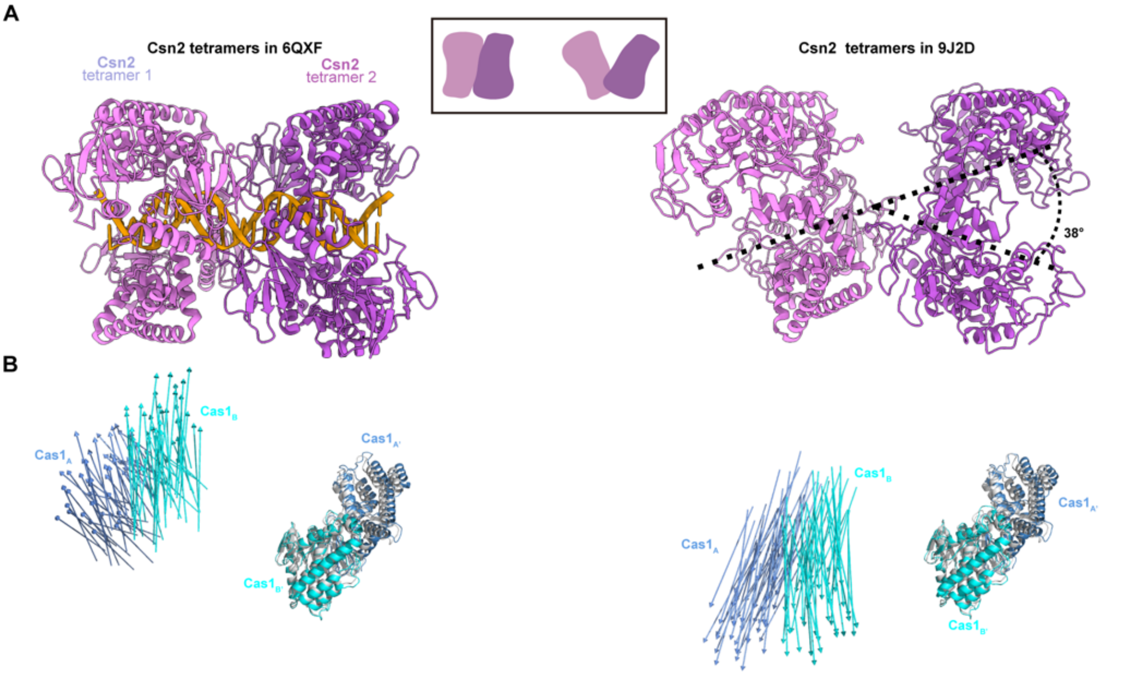


**Figure S4 Csn2 tilting controls the conformational change of Cas1$_4$-Cas2$_2$ in the Cas9/RNA$_2$-Csn2$_8$-Cas1$_8$-Cas2$_4$ supercomplex.**

**(A)** Comparison of Csn2 tetramers in different complex. The V-shape architecture with 38° tilting of two Csn2 tetramers in the Cas9/RNA$_2$-Csn2$_8$-Cas1$_8$-Cas2$_4$ supercomplex. The tilt angle of Csn2 is measured by aligning the central cavities of the two Csn2 tetramers.

**(B)** The vector map representation of Cas1 conformational change. One Cas1 dimer of the upper (left) or lower (right) Cas1$_4$-Cas2$_2$ is superimposed with protospacer-bound Cas1$_4$-Cas2$_2$ structure (PDB 5XVN, grey), and the resulting shift of the other Cas1 dimer is highlighted by vector map.

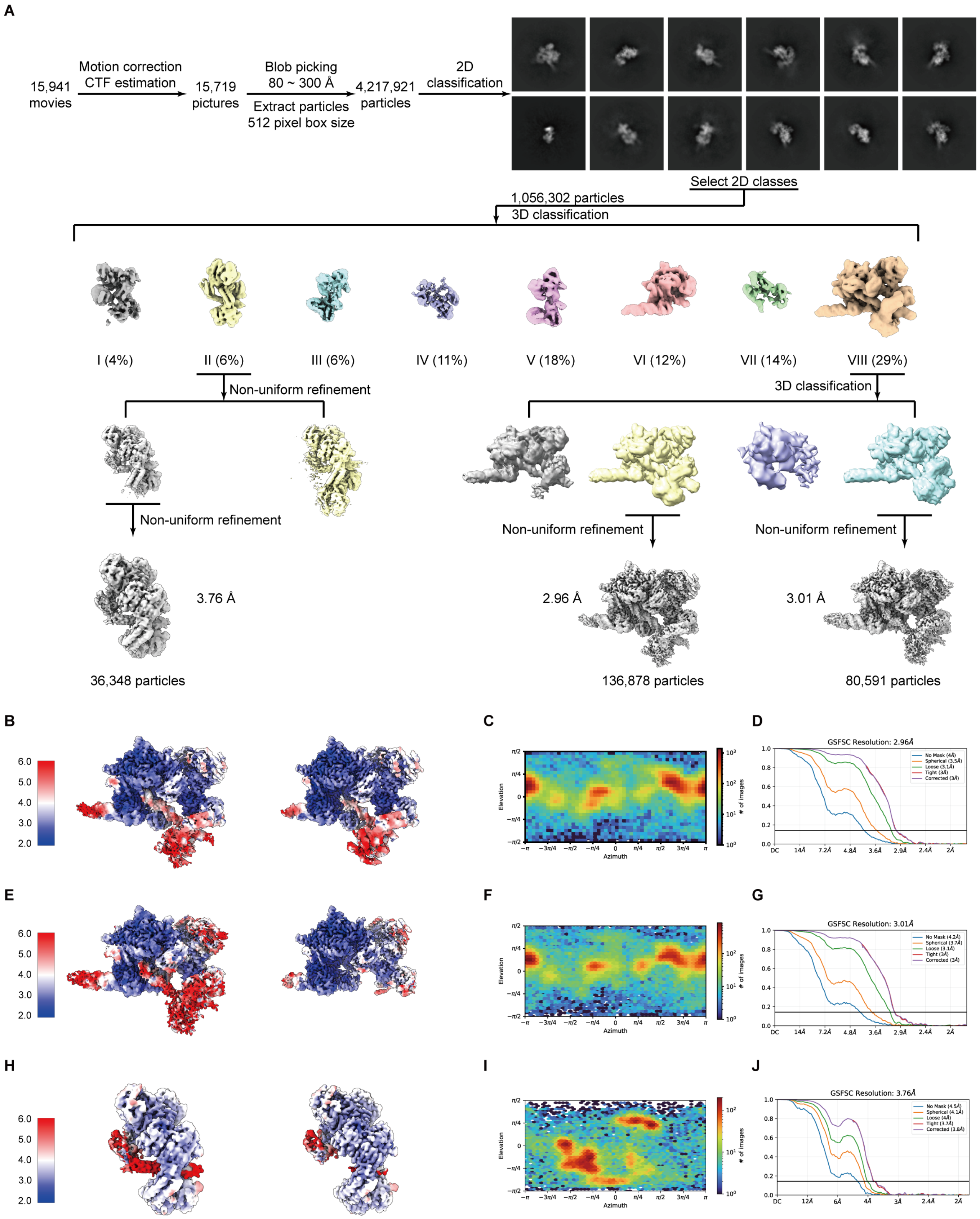


**Figure S5 Single-particle cryo-EM analysis of DNA binding complex**

**(A)** Cryo-EM reconstruction workflow for DNA binding complex.

**(B-D)** Cryo-EM density of the Prespacer catching complex (state I) colored by the local resolution (**B**). Left, the original cryo-EM map. Right, the cryo-EM maps filtered to their estimated overall resolution. The FSC curves (**C**) and the viewing direction distribution plot (**D**) for data processing.

**(E-G)** Cryo-EM density of the Prespacer catching complex (state II) colored by the local resolution (**E**). Left, the original cryo-EM map. Right, the cryo-EM maps filtered to their estimated overall resolution. The FSC curves (**F**) and the viewing direction distribution plot (**G**) for data processing.

**(H-J)** Cryo-EM density of the Pre-integration subcomplex colored by the local resolution (**H**). The FSC curves (**I**) and the viewing direction distribution plot (**J**) for data processing.

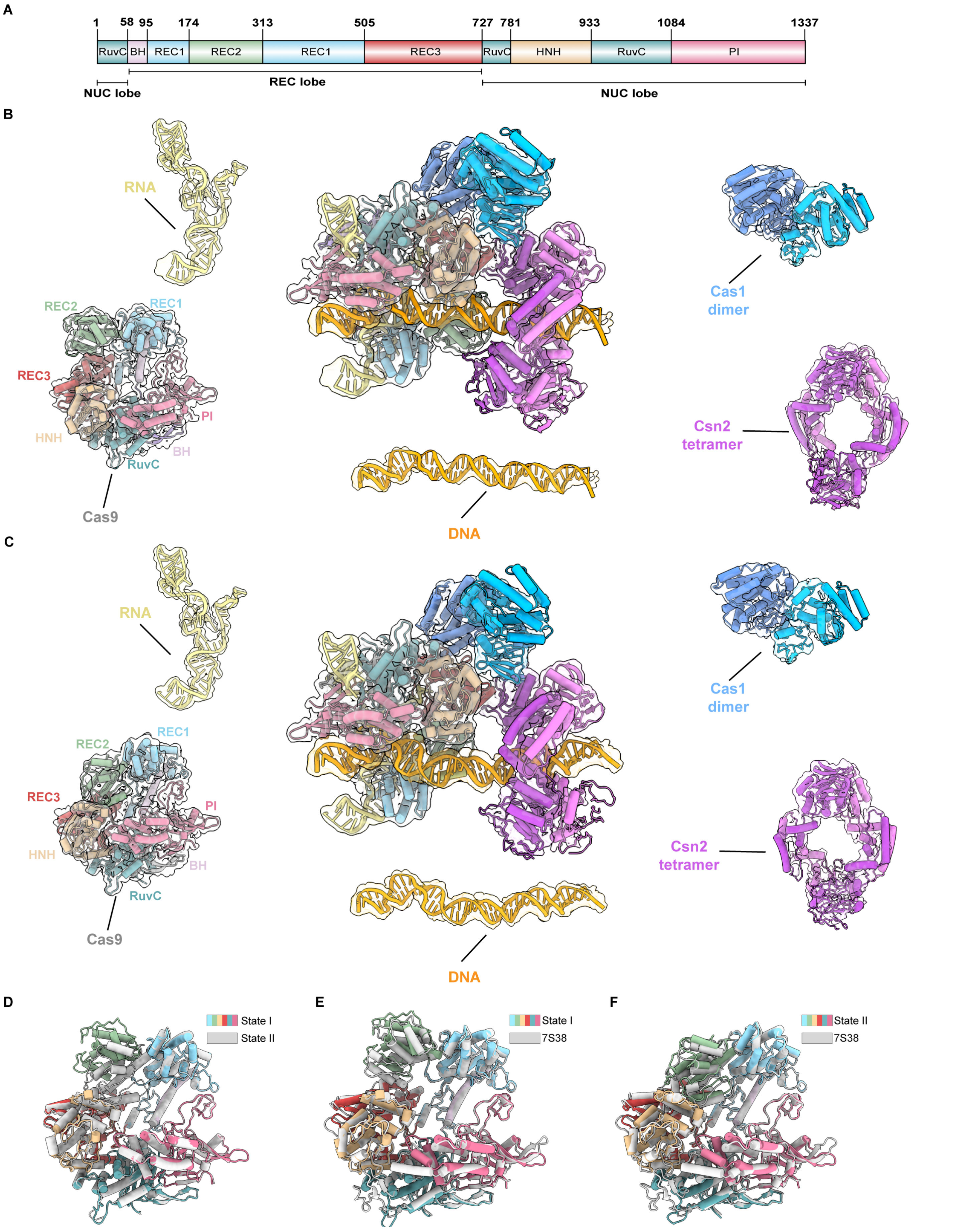
A
1
58
95
174
313
505
727
781
933
1084
1337
RuvC
BH
REC1
REC2
REC1
REC3
RuvC
HNH
RuvC
PI
NUC lobe
REC lobe
NUC lobe
B
RNA
REC2
REC1
REC3
PI
HNH
BH
RuvC
Cas9
DNA
Cas1
dimer
Csn2
tetramer
C
RNA
REC1
REC2
REC3
PI
HNH
BH
RuvC
Cas9
DNA
Cas1
dimer
Csn2
tetramer
D
State I
State II
E
State I
7S38
F
State II
7S38

**Figure S6 The cryo-EM structure of Cas9/RNA-Csn2-Cas1-DNA Prespacer catching complex**

**(A)** Domain organization of *Efa*Cas9.

**(B)** The structure model of prespacer catching complex state I shown in cartoon with filtered cryo-EM map shown as a transparent surface with contouring level in ChimeraX is 0.08. A Cas1 dimer (Blue and cyan), one Csn2 tetramer (purple and violet), RNA (yellow) and dsDNA (orange) can be docked well. Distinct domains of the Cas9 are resolved in the cryo-EM density.

**(C)** The structure model of prespacer catching complex state II shown in cartoon with filtered cryo-EM map shown as a transparent surface with contouring level in ChimeraX is 0.06. A Cas1 dimer (Blue and cyan), one Csn2 tetramer (purple and violet), RNA (yellow) and dsDNA (orange) can be docked well. Distinct domains of the Cas9 are resolved in the cryo-EM density.

**(D)** Cas9 structures in the two prespacer catching complex states exhibit distinct conformational changes within the REC2 domain, showing an RMSD of 0.649 Å over 1149 pruned atom pairs. and an RMSD of 6.455 Å over 1321Cα atoms.

**(E)** The Cas9 structure in the prespacer catching complex state I resembles SpyCas9/sgRNA/DNA with a 3 bp R-loop (PDB 7S38), showing an RMSD of 3.180 Å over 1306 Cα atoms.

**(F)** The Cas9 structure in the prespacer catching complex state II resembles SpyCas9/sgRNA/DNA lacking RNA:DNA base pairs (PDB 7S36), showing an RMSD of 2.482 Å over 1308 Cα atoms.

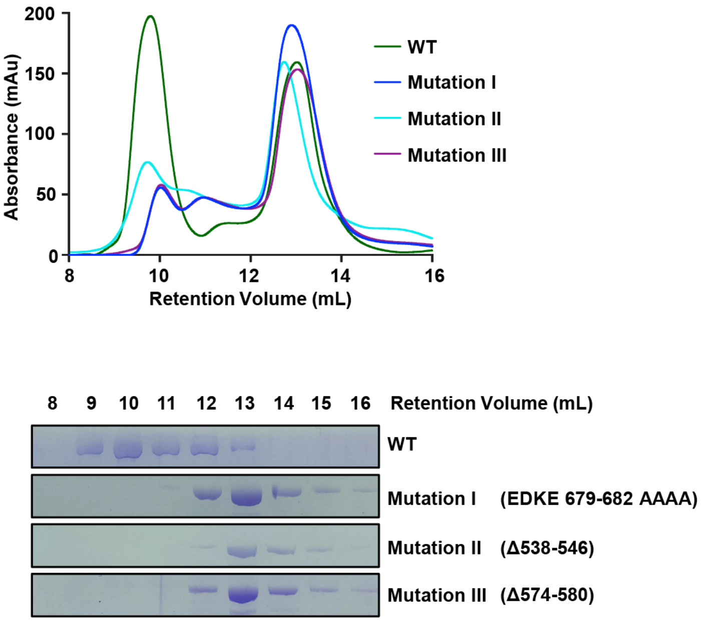


**Figure S7 Size exclusion chromatography analysis of Cas9 mutants.**

The interaction between the REC3 mutated Cas9 and Csn2-Cas1-Cas2 was assessed by size exclusion chromatography. The elution fractions were analyzed by SDS-PAGE.

.

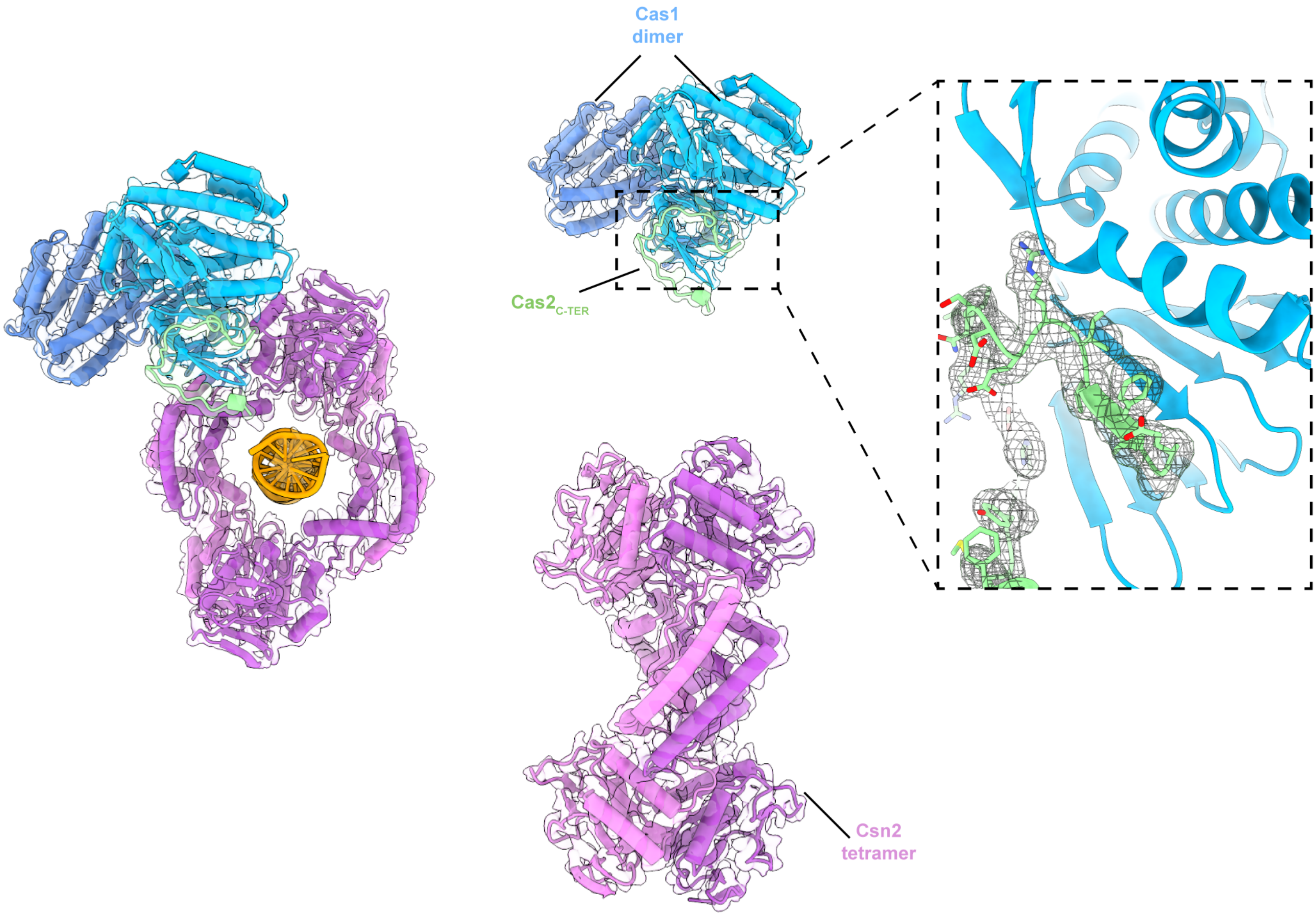


**Figure S8 The cryo-EM structure of Csn2-Cas1-Cas2 subcomplex.**

The structure model is shown in cartoon with sharpened cryo-EM shown as a transparent surface. A Cas1 dimer (Blue and cyan), a Csn2 tetramer (purple and violet) and dsDNA can be modeled well, but only one β-fold of Cas2 that directly interact with Cas1 can be built.

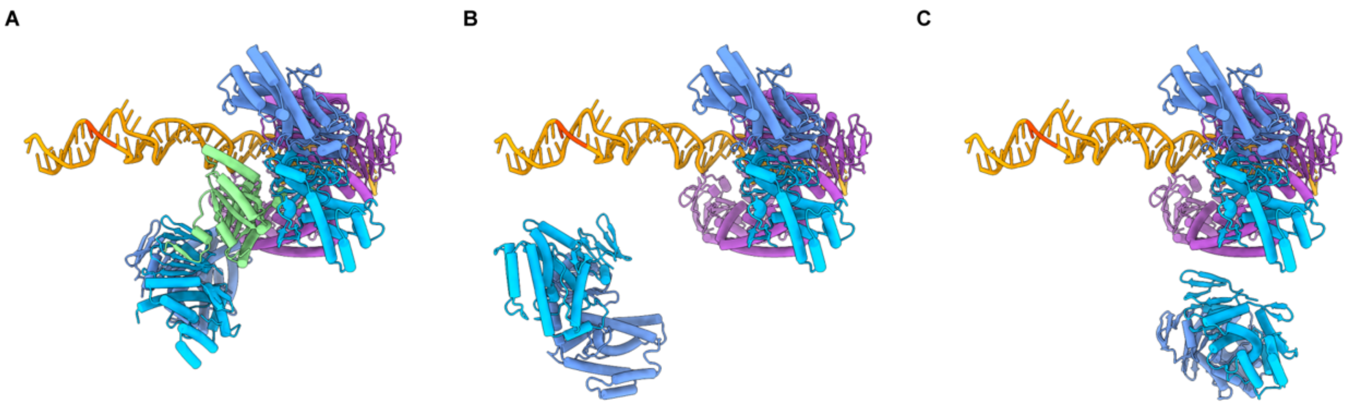


**Figure S9 Structural alignment of Cas1-Cas2 hexamers with the Cas1 dimer in the prespacer-catching complex .**

Alignment of Cas1-Cas2 hexamers from PDB 5XVN (**A**), the upper hexamer from the resting-state supercomplex (**B**), and the lower hexamer from the resting-state supercomplex (**C**) with the Cas1 dimer in the prespacer-capturing complex suggests DNA inaccessibility when Cas1 binds the Csn2 head domain.

**Video S1 Distinct Cas1-Csn2 interfaces in two different state complexs.**

Cas9 dissociation disrupts the REC3/Cas1/Csn2 interface, breaking the interaction between the Csn2 head domain and the Cas1 N-terminal domain. This destabilization induces Cas1 dimer sliding, forming a new interface between Cas1 and the hinge connecting the Csn2 head and tail domains.

**Table S1.**

Cryo-EM data collection, refinement and validation statistics.

| | Type II-A CRISPR integrase supercomplex, apo form (EMD-34441) (PDB 9J2D) | Type II-A CRISPR integrase prespacer catching complex, State I (EMD-61100) (PDB 9J2S) |
|---|---|---|
| **Data collection and processing** | | |
| Magnification | 130 k | 130 k |
| Voltage (kV) | 300 | 300 |
| Electron exposure (e–/Å$^2$) | 50 | 45 |
| Defocus range (μm) | -1.0 to -2.1 | -1.0 to -2.4 |
| Pixel Size (Å) | 1.100 | 0.932 |
| Symmetry imposed | C2 | C1 |
| Initial particle images (no.) | 542,743 | 1,0,56,302 |
| Final particle images (no.) | 52,810 | 136,878 |
| Map resolution (Å) | 3.90 | 2.96 |
| FSC threshold | 0.143 | 0.143 |
| **Refinement** | | |
| Initial model used (PDB code) | Alphafold3, 5XVN, 3C5U | Alphafold3, 9J2D, 3C5U |
| Model resolution (Å) | 3.6 | 3.4 |
| FSC threshold | 0.5 | 0.5 |
| Map sharpening *B* factor (Å$^2$) | DeepEMhancer | DeepEMhancer |
| Model composition | | |
| Non-hydrogen atoms | 36975 | 26405 |
| Protein residues | 4512 | 2771 |
| Nucleotide | 0 | 178 |
| Ligands | 12 | 0 |
| *B* factors (Å$^2$) | | |
| Protein | 105,85 | 115.13 |
| Nucleotide | 0 | 167.56 |
| Ligand | 95.70 | 0 |
| R.m.s. deviations | | |
| Bond lengths (Å) | 0.003 | 0.005 |
| Bond angles (°) | 0.605 | 0.814 |
| Validation | | |
| MolProbity score | 2.46 | 2.64 |
| Clashscore | 7.10 | 13.06 |
| Poor rotamers (%) | 5.38 | 4.74 |
| Ramachandran plot | | |
| Favored (%) | 91.07 | 91.58 |
| Allowed (%) | 8.64 | 8.09 |
| Disallowed (%) | 0.29 | 0.33 |

| | Type II-A CRISPR integrase prespacer catching complex, State II (EMD-61101) (PDB 9J2T) | Type II-A CRISPR integrase pre-integration subcomplex, (EMD-61097) (PDB 9J2I) |
|---|---|---|
| **Data collection and processing** | | |
| Magnification | 130 k | 130 k |
| Voltage (kV) | 300 | 300 |
| Electron exposure (e–/Å$^2$) | 45 | 45 |
| Defocus range (μm) | -1.0 to -2.4 | -1.0 to -2.4 |
| Pixel Size (Å) | 0.932 | 0.932 |
| Symmetry imposed | C1 | C1 |
| Initial particle images (no.) | 1,0,56,302 | 1,0,56,302 |
| Final particle images (no.) | 80,591 | 50,477 |
| Map resolution (Å) | 3.01 | 3.76 |
| FSC threshold | 0.143 | 0.143 |
| **Refinement** | | |
| Initial model used (PDB code) | Alphafold3, 9J2D, 3C5U | Alphafold3, 5XVN, 3C5U |
| Model resolution (Å) | 3.4 | 4.1 |
| FSC threshold | 0.5 | 0.5 |
| Map sharpening *B* factor (Å$^2$) | DeepEMhancer | DeepEMhancer |
| Model composition | | |
| Non-hydrogen atoms | 26415 | 13115 |
| Protein residues | 2770 | 1479 |
| Nucleotide | 180 | 50 |
| Ligands | 0 | 6 |
| *B* factors (Å$^2$) | | |
| Protein | 188.50 | 89.4 |
| Nucleotide | 150.49 | 354.92 |
| Ligand | 0 | 95.87 |
| R.m.s. deviations | | |
| Bond lengths (Å) | 0.005 | 0.005 |
| Bond angles (°) | 0.972 | 1.226 |
| Validation | | |
| MolProbity score | 2.50 | 2.17 |
| Clashscore | 11.54 | 7.00 |
| Poor rotamers (%) | 3.38 | 3.89 |
| Ramachandran plot | | |
| Favored (%) | 91.10 | 95.15 |
| Allowed (%) | 8.68 | 4.37 |
| Disallowed (%) | 0.22 | 0.48 |
